# Direct cryo-ET detection of native SNARE and Munc13 protein bridges using AI classification and preprocessing

**DOI:** 10.1101/2024.12.18.629213

**Authors:** Daniel H. Orozco-Borunda, Christos Papantoniou, Nils Brose, Vladan Lučić

**Affiliations:** Department of Molecular Structural Biology, Max Planck Institute of Biochemistry, Am Klopferspitz 18, 82152 Martinsried, Germany; Department of Molecular Neurobiology, Max Planck Institute of Multidisciplinary Sciences, City Campus, 37075 Göttingen, Germany; iHuman Institute, ShanghaiTech University, 393 Middle Huaxia Road, Pudong, Shanghai, China

## Abstract

Synaptic transmission requires Munc13 and SNARE proteins for synaptic vesicle priming and fusion. Cryo-electron tomography detected multiple types of Munc13- or SNARE-dependent dependent molecular bridges that tether synaptic vesicles to the presynaptic active zone plasma membrane. To integrate the molecular scenario with structural observations, we obtained de novo, in situ cryo-electron tomography averages of native, mammalian SNARE- and Munc13-dependent tethers. These provide direct evidence that Munc13 and a complex comprising SNARE proteins link synaptic vesicles to the active zone membrane. Furthermore, we determined the plausibility of different molecular compositions of tethers, placed constraints on their conformations and positioning, and proposed the existence of a complex downstream of Munc13 and upstream of SNARE complex formation. Because the detection and subtomogram averaging of membrane-bridging complexes is complicated by the presence of two lipid membranes and multiple protein species and conformations, we developed preprocessing methods and feature-based AI classifiers that outperformed standard methods.

## Introduction

Synaptic transmission comprises a number of precisely organized biochemical events. At the presynaptic terminal, Munc13-1 is necessary for synaptic vesicle (SV) priming, a process that renders individual SVs capable of fusion with the presynaptic membrane upon Ca^2+^ influx into the presynaptic terminal [1, 2]. Munc13 proteins and Munc18-1 facilitate the formation of the synaptic soluble N-ethylmaleimide-sensitive factor attachment protein receptor (SNARE) complex comprising SNAP25, syntaxin-1 and synaptobrevin-2 [3], and the full assembly of the SNARE four-helix bundle drives the fusion of primed SVs and subsequent neurotransmitter release [4, 5]. Synaptotagmin-1 is the major Ca^2+^sensor for synchronous SV fusion, acting by binding Ca^2+^ via two C_2_ domains (C_2_A and C_2_B), while complexins regulate SNARE complex function [6, 7, 8, 9]. Synaptotagmin-1 and complexin-1 bind distinct sites of the SNARE complex and are thought to bind simultaneously during SV fusion [10, 11, 12].

Munc13-1 is thought to execute its SNARE-regulating priming function via its MUN-domain by regulating the conformation of the SNARE protein syntaxin, and is stimulated by Ca^2+^- calmodulin binding to an amphipathic helical motif, by diacylglycerol (DAG) binding to a C_1_domain, and by Ca^2+^-phospholipid binding to the central C_2_B domain [13, 14, 15, 16, 17, 18]. Data from reconstituted systems showed that efficient liposome fusion requires the C_1_, C_2_B, MUN and C_2_C domains of Munc13-1, and that Munc13-1 and the SNARE complex can simultaneously bind two membranes [19, 20, 21, 3]. Together, these data led to various models of SNARE complex and Munc13 architecture and positioning with respect to SVs and the plasma membrane [19, 22]. However, there is no direct evidence that Munc13 and the SNARE complex link SV and plasma membranes in situ.

The function of membrane-bound complexes in synaptic transmission and other cellular processes depends to a large extent on their native state and cellular environment. Cryo-electron tomography (cryo-ET) provides a molecular-level visualization of the presynaptic terminal [23] and is uniquely suited for label-free imaging of protein complexes at single nanometer resolution in situ, i.e. in their native composition, conformation, and protein and lipid environment [24, 25]. Subtomogram averaging reduces noise and increases interpretable information by aligning and averaging subtomograms of individual complexes (termed particles) [26, 27]. Although averages of small proteins have been obtained, such as 220 kDa Arp 2/3 in membrane-extracted and chemically fixed samples, and ∼200 kDa ribosome-free OST complex on isolated microsomal membranes [28, 29], in situ averaging of non-periodic, sparsely distributed small proteins is severely limited because of difficulties in their detection and alignment.

Previous molecular-level cryo-ET studies of the presynaptic terminal showed that pleomorphic protein bridges tether SVs to the presynaptic plasma membrane at the active zone, and are crucial for neurotransmitter release [30, 31, 23]. To avoid the present nomenclature ambiguity, we here define tethers in the structural sense, as all directly observed bridges linking SVs with the plasma membrane because the molecular identity of proteins visualized by cryo-ET is not known a priori [30, 32]. More recently, combining genetic manipulations with cryo-ET, we showed that tethers shorter than 12 nm (intermediate tethers) depend on the presence of Munc13 and are responsible for bringing SVs closer than 10 nm to the plasma membrane, while tethers shorter than 6 nm depend on the presence of the SNARE protein SNAP25 and bring SVs closer than 5 nm to the active zone [33]. We also found that many tethers were large enough to accommodate SNARE complex or Munc13, which was recently confirmed [34].

To determine molecular identity and conformation of protein complexes forming tethers between SVs and the active zone membrane, we obtained de novo, in situ subtomogram averages of short SNARE dependent and intermediate Munc13 dependent tethers and fit the available atomic models into these densities. To alleviate the difficulties caused by the presence of two lipid membranes, located at a variable distance to each other and one highly curved, and by the molecular heterogeneity of tethers and the crowded cellular environment, it was necessary to devise new subtomogram averaging methods, as well as artificial intelligence (AI) based classification methods. Our data show that Munc13-1 and a complex comprising SNARE proteins link SVs to the active zone plasma membrane.

## Results

### Subtomogram processing developments for membrane-bridging complexes in situ

Previously, we acquired a number of cryo-electron tomograms of rodent central nervous system synapses, segmented SVs and plasma membranes (together called boundaries), and used the hierarchical connectivity procedure to detect, segment and characterize tethers within presynaptic terminals (Figure 1A, B) [35, 33].

**Figure 1:**
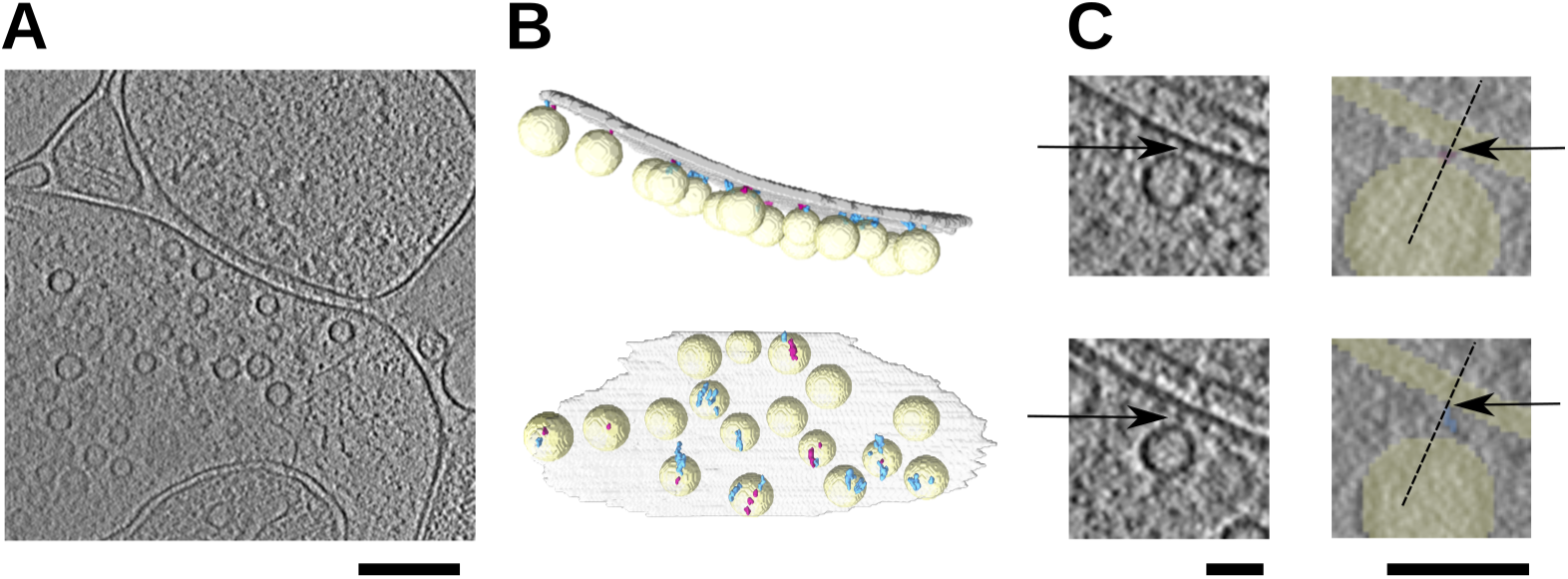
Preprocessing developments. (A) Tomographic slice of a synapse. Slice thickness 5.27 nm, Gaussian low-pass filtered at 0.7 pixels, scale bar 200 nm. (B) 3D visualization of the same presynaptic terminal showing short tethers (red) and intermediate tethers (blue), views from the side and the postsynapse. (C) Short (upper row) and intermediate tether from the same tomogram (lower row) shown as magnified insets from A (slice thickness 0.88 nm, without filtering, arrows point at tethers). The right column slices are overlaid with segmented components. Dashed lines show the direction perpendicular to the membrane and arrows point at the contact between tethers and the plasma membrane. Scale bars 40 nm. Plasma membrane (grey in B, yellow in C), SVs (yellow).

Here we determined the coordinates of tether contacts with the plasma membranes and the membrane orientation (Figures 1C, S1), and extracted subtomograms of tethers (particles), which constitute the “Plain” tether set (Figure S2B). In addition, we extracted subtomograms from boundaries at the same locations as the tether particles, resulting in the “boundaries” particle set (Figure S2B). Thus, neither visual detection, nor external templates were used to determine tether positions.

Because the small size of tethers and the presence of membranes negatively affect tether averaging, we devised the following methods to improve subtomogram alignment and averaging. First, we preprocessed tether particles by selectively low-pass filtering regions of tether particles corresponding to the boundaries (plasma membrane or SVs) to obtain“filtered” datasets (Figures S2A, S1). In this way, the alignment of membranes, which is based on lower resolution information, can constrain the particle alignment parameter space, while the higher resolution information pertaining to tethers can provide a more precise alignment, thus preventing alignment of tethers to membranes and other proteins located in their vicinity. Furthermore, we used the segmented tethers to generate particle-specific masks focused on tethers, resulting in the “focused” tether particle sets.

Second, we used the location of the plasma membrane to set the initial tether alignment parameters so that the z-axis was perpendicular to the plasma membrane and the cytosolic membrane faces were aligned with each other. This made it possible to use constrained subtomogram alignment, whereby only limited changes from the initial values are allowed for the position and two Euler angles, while the third angle is not constrained [29].

Third, we averaged the boundaries particle set using the final alignment parameters obtained from the tether averaging. Because the resulting boundaries average was aligned with the tether average, we used it to separate the cytosolic part of the tether average.

In sum, we designed a set of particle preprocessing approaches and devised subtomogram averaging improvements to focus the alignment on tethers, to reduce the influence of lipid membranes and the surrounding proteins, and to constrain the space of the alignment parameters that is explored during subtomogram averaging. This procedure yields de novo averages because no external reference is used at any point.

### Subtomogram averaging of short tethers (SNARE complexes)

We employed the methods described above to generate particle sets corresponding to different preprocessing approaches and to determine de novo average densities of short (SNARE-dependent) tethers.

A visual assessment of the averages provided estimates of optimal filter cutoff frequencies, and indicated that it is advantageous to filter SVs more strongly than the plasma membrane, dilate the particle-specific masks over cytosol only and not over the membranes, and combine focusing with filtering. This led us to the following four preprocessing approaches for tether particle sets: (i) Plain, (ii) low-pass filtered SVs (Filtered SV), (iii) low pass filtered SVs and plasma membrane (Filtered SV&PM) and (iv) tether focused (Focused) (Figure S2).

All subtomogram averages obtained from the four short tethers particle sets defined above contained a well-defined density comprising a bridge between a SV and the plasma membrane and a laterally extended density, consistent with the expected basic morphology of SNARE-dependent tethers (Figure S3A). The Plain tether set average differed from the rest in that the lateral density was slanted, as opposed to parallel to the plasma membrane in the other three averages, with a further distinction that the Plain average density contained relatively large, disconnected pieces. The lateral density was closer to the SV in the Filtered SV and Filtered SV&PM averages than in the Focused one.

In all cases, the rotation angles around the direction normal to the plasma membrane were well-distributed within individual synapses (Fig. S4A). As expected, the other two angles depend on the orientation of the synapse, together supporting the validity of our methods. The resolution determined by the “gold standard” (as implemented in Relion) was in the range of 2.8-3.3 nm (Figure S4C).

We also obtained the boundaries set averages corresponding to all tether averages (Fig. S4B), and used them to generated membrane masks and to extract the cytosolic parts of tethers. Density traces calculated from boundary averages along lines perpendicular to the plasma membrane confirmed that the membranes were well resolved and showed that the shortest distances between the SV and the plasma membrane were the same for all four preprocessing cases, together supporting the validity of the methods used (Figure S4D). Applying 3D classification resulted in inferior averages, arguing against the existence of distinct subsets of our particle sets.

Among the previously proposed SNARE complex atomic models that include synaptotagmin C_2_B domain and complexin-1, the model that has the strongest biochemical evidence states that synaptotagmin C_2_B domain binds SNARE helices via the so-called primary interface, and can bind to the plasma membrane (PIP2) via Ca^2+^-dependent and Ca^2+^-independent mechanisms via the polybasic region (Figure S3B) [22, 36]. In this model, complexin-1 and the synaptotagmin C_2_B domain are located on opposite sides of SNARE helices. According to the tripartite model, synaptotagmin C_2_B domain binds SNARE helices and complexin-1 via the tripartite interface, thus positioning complexin-1 and the synaptotagmin C_2_B on the same side of SNARE helices [12]. Finally, a model was proposed where two synaptotagmin C_2_B domains bind SNARE helices, one via the primary and the other via the tripartite interface. We combined these models with the model that contained the helical part of complexin-1, resulting in three models that all contained the SNARE four-helix bundle, complexin-1 and one or two synaptotagmin C_2_B domains [12].

To investigate the plausibility of the proposed bindings, we fit the three SNARE complex models into the average densities by manual rigid-body fitting. The Filtered SV, Filtered SV&PM and Focused averages contained a laterally extended cylindrical region close to the SV and an attached globular region towards plasma membrane, which could accommodate almost the entire SNARE four-helix bundle and the synaptotagmin C_2_B domain (Figs 2 S3A). In contrast, because of the slanted orientation of the lateral cylindrical density in the Plain average, the N- and C-termini of the SNARE four-helix bundle had to be positioned very close to the SV and plasma membrane, which casts doubt on the validity of this average.

**Figure 2:**
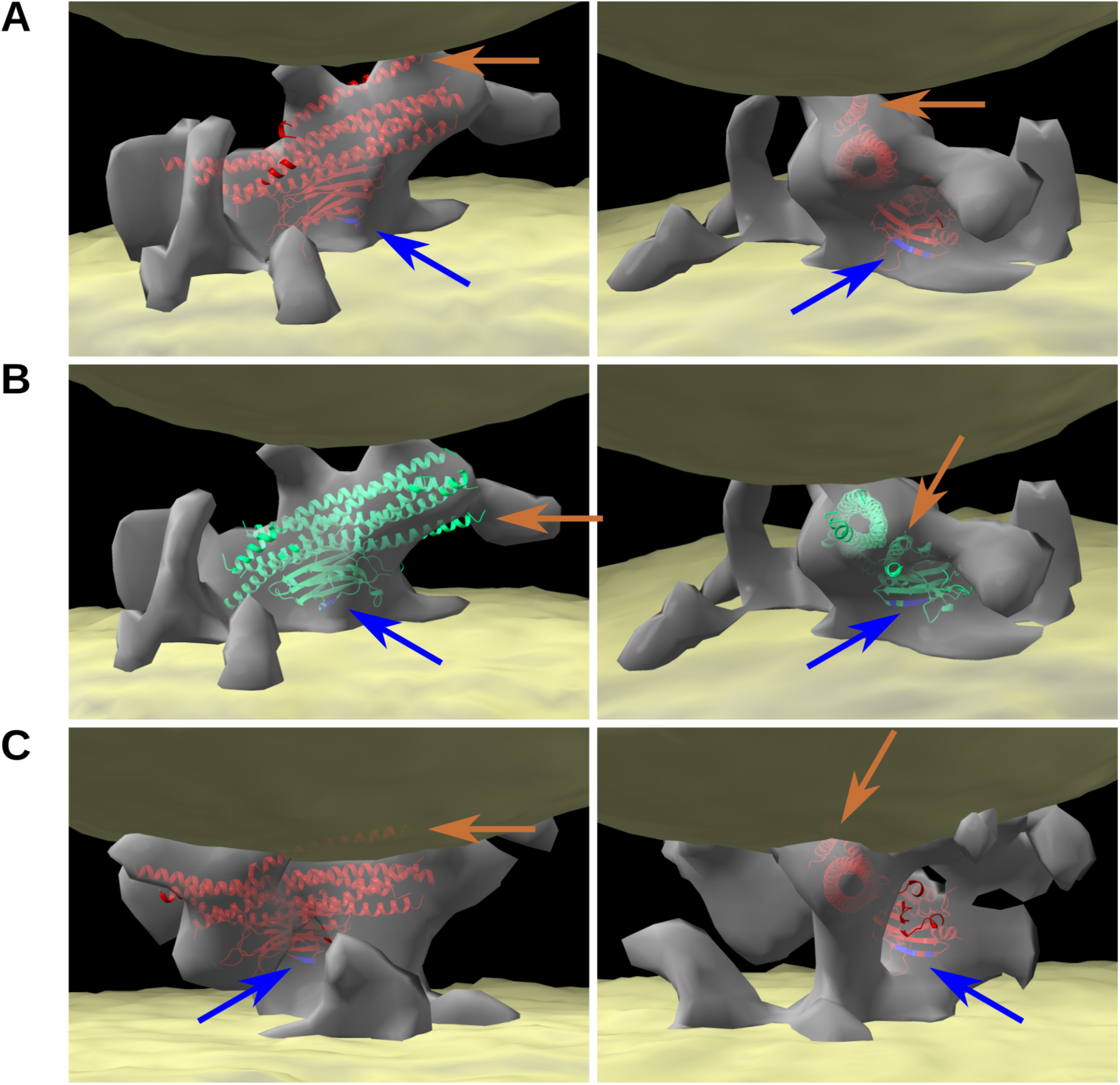
Short tether (SNARE complex) de novo average densities. (A) Focused preprocessing, primary atomic model. (B) Focused preprocessing, tripartite atomic model. (C) Filtered SV&PM preprocessing, primary atomic model. Average tether density is shown in grey and average boundaries comprising SV (above) and plasma membrane (below) in yellow. Images in the left column have the SNARE N-terminals on the left, while the right column shows the view from the SNARE C-terminals. Blue arrows point to synaptotagmin C_2_B polybasic region, brown arrows to complexin-1. All images are drawn to scale.

In both Filtered averages, the SNARE four-helix bundle had to be placed closer to the SV than to the plasma membrane, while in the Focused average the SNARE four-helix bundle was in the middle, the latter providing the optimal positioning of the polybasic region of synaptotagmin C_2_B domain, close to the plasma membrane (Figure 2). This positioning of the SNARE helices caused complexin-1 to be entirely occluded by the SV in both Filtered averages, while in the Focused average, most of complexin-1 fit into the density and it was located completely outside the SV. In all these cases, the model did not account for the entire density, implying that the averages could accommodate additional proteins.

Importantly, fitting of all models was constrained by the requirement that the polybasic residues of the synaptotagmin C_2_B domain face the plasma membrane, except when the polybasic residues were oriented towards the SV (Figs 2 S3A).

Like the primary model, the tripartite model with the C_2_B domain oriented towards the SV provided good fits into the the Filtered and the Focused average densities (Figure S3). The Plain average could not accommodate the tripartite model because a large part of the C_2_B domain collided with the plasma membrane, in addition to the slanted orientation of the lateral density, causing the same problem as for the primary model fit. Most of complexin-1 did not fit into the two Filtered averages, while it almost completely fit into the Focused average. Like for the primary model, the polybasic residues of the synaptotagmin C_2_B domain were positioned closer to the plasma membrane for the Focused than for the two Filtered averages.

When the tripartite model was oriented so that the synaptotagmin C_2_B domain faced the SV, it was not possible to fit it into any of the four averages because either fitting the synaptotagmin C_2_B domain into the densities close to the SV did not leave a sufficient space to fit the SNARE four-helix bundle, or the C_2_B domain could not be fit. (Figure S3). Similarly, the two synaptotagmin C_2_B domain SNARE model could not fit into any of the averages.

Together, the short tether averages obtained from different preprocessing approaches are best compatible with the notion that short tethers are composed of SNARE protreins, synaptotagmin-1 and complexin-1 (132 kDa in total). Furthermore, our results argue against positioning the synaptotagmin C_2_B domain close to the SV. Both the primary and the tripartite model where the synaptotagmin C_2_B domain is located close to the plasma membrane provided equally good fits, the main difference being the positioning of complexin-1. Concerning methodology, we successfully optimized the particle preprocessing and showed that focusing on the dilated tether segments and differential low-pass filtering of membranes provided crucial subtomogram averaging improvements.

### Annotation and feature selection

Previously, we showed that Munc13 is required for the intermediate tethers (6-12 nm in length), and that many of them were too small to accommodate the atomic model of Munc13 [33]. Because these results were obtained by manual fitting into individual tethers, a labor intensive and error-prone procedure, we proceeded to develop artificial intelligence (AI) based supervised classifiers that can predict which tethers can accommodate the Munc13 model.

We annotated an additional set of tethers by assigning them into three classes: the first comprising tethers that could accommodate the Munc13 model [37] (Munc13 class), the second where only the MUN domain but not the entire Munc13 model could fit (MUN class) and the third that could not even accommodate the MUN domain (NoFit class).

To generate features to be used for classification, we used morphological, greyscale density and topological properties of tethers from our previous tether characterization [33] and further analyzed the, resulting in 14 Basic features. Furthermore, to make features more suitable for AI classification, we expanded the feature set by combining the Basic features and selected optimal features by removing features that were highly correlated or had low univariate ANOVA and Mutual information scores, resulting in 16 features termed the Selected set (Figure S5). Their distributions were normalized and highly skewed features were logarithmically transformed, resulting in robust, standard, logrobust and logstd scaled feature sets (Figure S6).

### Classification of intermediate (Munc13-dependent) tethers by machine learning (excluding deep learning)

To build a machine learning classifier, it is necessary to choose an appropriate algorithm (model type) and an evaluation metric, and train and evaluate the model.

We chose the macro variant of the F1 evaluation metric, which gives the same weight to each class, because our dataset was imbalanced and we did not want the classification to be dominated by the largest class. Nevertheless, we also calculated accuracy and weighted F1 score, which applies higher weights to classes with more elements.

We first considered Logistic regression model (LogReg) with L2 (Tikhonov) regularization. To perform unbiased optimization, we randomly divided the annotated data into training and test sets, employed cross-validation for hyperparameter search on the training set, and evaluated the model on the test set. Because the separation into the training and the test sets may have an influence on the results for small datasets, the cross-validation was repeated for multiple random separations. Overall, test set scores were higher for the Selected than for the Basic test set, and higher for the logrobust and logstd than for the robust scaling (Table S1), The best F_1_macro score 0.75 ± 0.03 (mean ± std, N=22) was obtained for the Selected features and logstd scaling. Repeating cross-validation with volume-stratification did not improve the scores (F_1_macro 0.73 ± 0.03 for the Selected features and logstd scaling), indicating that the observed score variability over data separations was not caused by different distribution of tether volume in the training and test sets.

The training scores were somewhat higher than the testing scores, indicating that there was some degree of overfitting (Table S1). Testing and training scores for the Selected feature set over a range of hyperparameter *C* values showed that the optimal performance was obtained for *C* = 1, because for lower values the (test) scores were lower, while for larger values the model overfitted (Figure S7 A). Furthermore, the F_1_ macro test scores remained below 1 for all values of *C* (Table S1, Figure S7 A), indicating that the linear regression model is not powerful enough to fit the data. Therefore, we augmented the Selected feature set with all order 2 polynomial combinations of the features. Although the model was powerful enough to fit the training data perfectly (for *C >* 100), it showed large overfitting, arguing against this approach (Figure S7 B).

The support vector machine (SVM) classifier with Gaussian kernel allows non-linear classification, in contrast to LogReg. To investigate this model, we followed the same approach that we used for LogReg. Feature selection and scaling resulted in minimal changes and volume stratification did not improve the scores. The best F_1_macro score was 0.73 ± 0.06 (Table S1) and the optimal performance was obtained for hyperparameter values *C* = 100 and *γ* = 0.001 (Figure S7C). For some hyperparameter values, this model was able to fit the training data perfectly while strongly overfitting, showing that although being more powerful than LogReg, it did not provide a better classification.

Among the other model types that we applied to the tether classification, Decision trees and Gradient boosted trees show very high overfitting, while Ada boosted LogReg showed F_1_ macro scores below 0.6.

In sum, we identified two best models and showed that feature design, selection and scaling improved the evaluation scores.

### Classification of intermediate (Munc13-dependent) tethers by deep learning

We implemented a feature-based artificial neural networks (NN) classifier comprising a stack of fully connected layers and used the features defined above as the network input. The search for optimal hyperparameter values and feature sets was carried out by a series of cross-validations on the training set, and the trained models were evaluated on the test set. Because NNs have a high number of hyperparameters, initial searches involved simultaneous variation of multiple hyperparameters, while the final optimization stage contained a series of single hyperparameter optimizations. Model performance was evaluated using custom implemented F_1_ macro score, and we also calculated accuracy and F_1_ weighted score.

The scores for the best network were accuracy 0.90, F_1_ macro 0.82 and F_1_ weighted 0.90, while the training scores were a little higher, showing some overfitting (Figure S7D). The optimal architecture comprised four fully connected 18-node hidden layers. Reducing or increasing the number of 18-node layers, and changing the number of layers while keeping the total number of nodes constant resulted in significantly lower F_1_ macro scores (0.59 - 0.74). Furthermore, decreasing the number of layers resulted in larger overfits, while increasing the number led to a larger variability between training epochs.

The Selected set of features, with standard scaling, provided the highest score. Other combinations did not show clear tendencies, indicating that the interplay between feature selection and scaling plays an important, but complex role. Among other hyperparameters, the choice of the optimizer and the dropout rate were the most important. F_1_ macro loss function provided better results than cross-entropy during the cross-validation, but the scores for the optimal network were essentially the same.

In sum, extensive optimization allowed us to develop a deep neural net classifier that surpasses the performance of the machine learning algorithms investigated in the previous section.

### Subtomogram averaging of intermediate (Munc13-dependent) tethers

Intermediate tethers were classified by LogReg, SVN and NN classifiers that were trained and optimized as described above. Each classification yielded three classes (Munc13, MUN and NoFit). In addition, we classified all intermediate tethers using the conventional image-based 3D classification (Conventional3D). We then obtained de novo subtomogram averages for all classification, class and preprocessing combinations, resulting in 48 averages. In all cases, we followed the processing improvements described above for short tethers, obtained boundaries averages and extracted cytosolic parts of all intermediate tether class averages.

All 16 Munc13 class averages showed densities comprising a short stem attached to the plasma membrane and a longer branch extended at shallow angles towards the SV (Figure 3, S8A). Shorter branches were also present, indicating that other proteins may be present in the vicinity of intermediate tethers, except in the averages obtained from the Focused datasets. Plasma membrane regions were well defined, indicating a good alignment of individual subtomograms.

**Figure 3:**
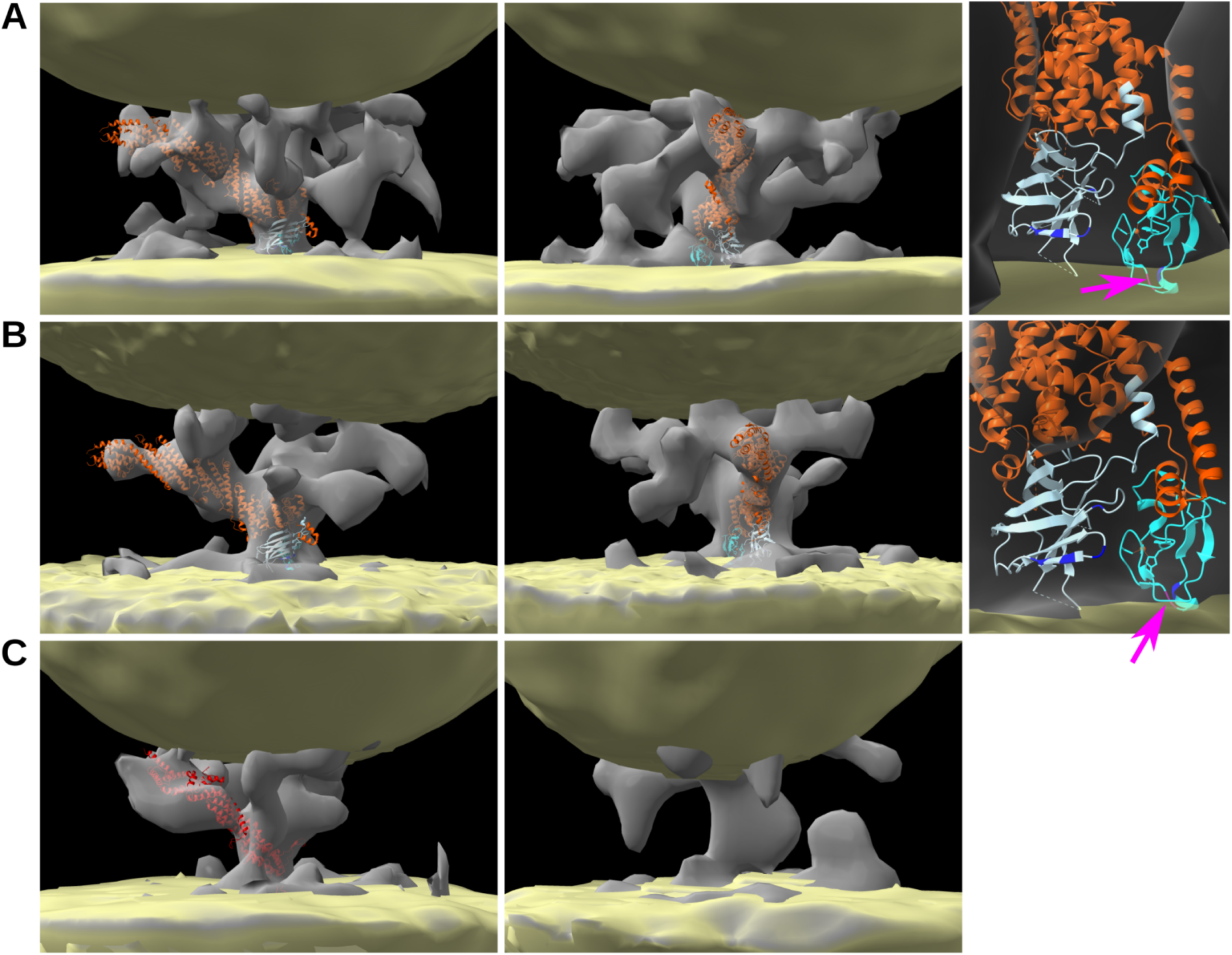
Intermediate tether (Munc13-dependent) de novo average densities. (A) Munc13 class average from NN classification, Filtered SV&PM preprocessing. (B) Munc13 class average from LogReg classification, Filtered SV preprocessing. (A) and (B) Panels on the right show the polybasic region of C_1_ and C_2_B domains (blue) and the DAG binding site (magenta, arrows). (C) MUN and NoFit class averages, both from NN classification, Filtered SV&PM preprocessing. Average tether density is shown in grey and average boundaries comprising SV (above) and plasma membrane (below) in yellow.

Visual assessment of all Munc13 class averages indicate that the three AI classifiers performed better than the Conventional3D classification (Figure S8A). Furthermore, Conventional3D classification resulted in the smallest number of particles in the Munc13 class compared with other classifications.

In all cases, like for short tethers, the final particle rotation angles around the plasma membrane normals were well-distributed within individual synapses (Fig S9A), supporting our methods. The resolutions determined by the “gold standard” were in the range of 3.3-3.5 nm, except for the Conventional3D Plain (3.7), Focused Plain (4.3), and NN Plain (4.3) (Figure S9B).

Density traces obtained from boundaries averages showed a higher slope (slower rate of change) at the SV membrane than at the cytosolic face of plasma membrane, which means that the SV surface was less well defined than the plasma membrane surface (Figure S10A), consistent with the visually well resolved plasma membrane and somewhat fuzzy SV appearance. Together with the flattened appearance of the SV surface in the region facing the plasma membrane, our results show a variability in the localization of SVs, both in the directions perpendicular and parallel to the plasma membrane. This indicates that even within the same class, tethers do not assume the same orientation with respect to SVs.

Currently the most complete atomic model of Munc13 comprises the C_1_, C_2_B and MUN domains, covering 55% of the entire Munc13 sequence [37] (Figure S8B). It contains a large, rod-shaped MUN domain that is flanked by the lipid-binding C_1_-C_2_B and C_2_C domains, which are expected to bind the plasma and the SV membranes, respectively. In all classification and preprocessing cases, it was possible to fit and orient the Munc13 model so that the DAG binding site was in a direct contact and the polybasic region of C_1_ and C_2_B domains very close to the plasma membrane, while keeping most of the structure inside the density. The MUN domain fit into the long branch of the density and the rest of the Munc13 model in the stem, thus positioning the MUN domain C-terminal close to the SV (Figures 3, S8). The angle between the Munc13 model and the plasma membrane was in all cases smaller than 45°. Furthermore, the MUN domain did not fit well into smaller density branches, showing that our fits were unique, apart from small orientation modifications. Therefore, our data indicate that the average densities were consistent across preprocessing and classification methods, and also selective for the Munc13 model.

Overall, the density obtained by NN classification and Filtered SV&PM preprocessing provided the best fit, followed by the LogReg classification with Filtered SV average, as judged visually (Figure 3A, B). However, the fits were not perfect, as there were always residues that did not fit in the density, raising the possibility that Munc13 adopts a slightly different conformation in situ.

All MUN class averages obtained from NN classification showed a slanted branch-like density arising from the plasma membrane, which was sufficiently long to fit the MUN domain, but not the Munc13 model (Figure 3C, S11). They were very different from NN classification NoFit class averages, which were smaller, comprising a stubby main part perpendicular to the plasma membrane. This is in contrast to averages from the Conventional3D classification, where averages from the MUN and the NoFit classes for all preprocessing cases were fairly branched, had a similar size and could all accommodate MUN domain, even though the larger class averages were assigned to the MUN and the smaller to the NoFit class.

Furthermore, considering all preprocessing cases, NN classification traces showed a larger separation between classes than the Conventional3D classification (Figure S10B), indicating that NN classification yielded classes that were better resolved than the Conventional3D classification.

In sum, our data from the Munc13 class, from four different preprocessing approaches and four classification methods show that about two thirds of all intermediate tethers are morphologically consistent with the Munc13 model, and that they make an angle smaller than 45° with the plasma membrane. Furthermore, the NN classification data from MUN and NoFit classes indicate that a significant number of intermediate tethers are not composed of Munc13 and have a rod-like or stubby morphology. Regarding methodology, the development of the preprocessing approaches and the AI classification methods substantially improved subtomogram classification and averaging, and was crucial to reach the present conclusions. The NN classifier correctly learned to distinguish classes defined in the annotated data, and provided better resolved classes than the Conventional3D classifier.

## Discussion

We obtained de novo average densities of endogenous, mammalian short and intermediate tethers that represent protein bridges between SVs and active zone of presynaptic plasma membranes, and that had previously been determined to require the SNARE complex and Munc13 proteins, respectively [33]. Our data support the notion that these tethers represent Munc13, a complex comprising SNARE proteins, complexin-1 and synptotagmin-1, and one or two as yet unidentified complexes that may contain some of the SNARE proteins and require but are not composed of Munc13, at least not in an extended conformation like the one obtained previously [37]. Mapping these averages on a tomogram provides a unified view of the location, spatial orientation and the composition of the complexes that form tethers (Figure 4). The protein tethers were imaged in a fully hydrated, vitrified state by cryo-ET in situ, in their native composition, conformation, and protein and lipid environment. The structural information we obtained was sufficient to discriminate between currently available atomic models, which allowed us to determine the plausibility of different molecular compositions, and place constraints on their conformations and positioning.

**Figure 4:**
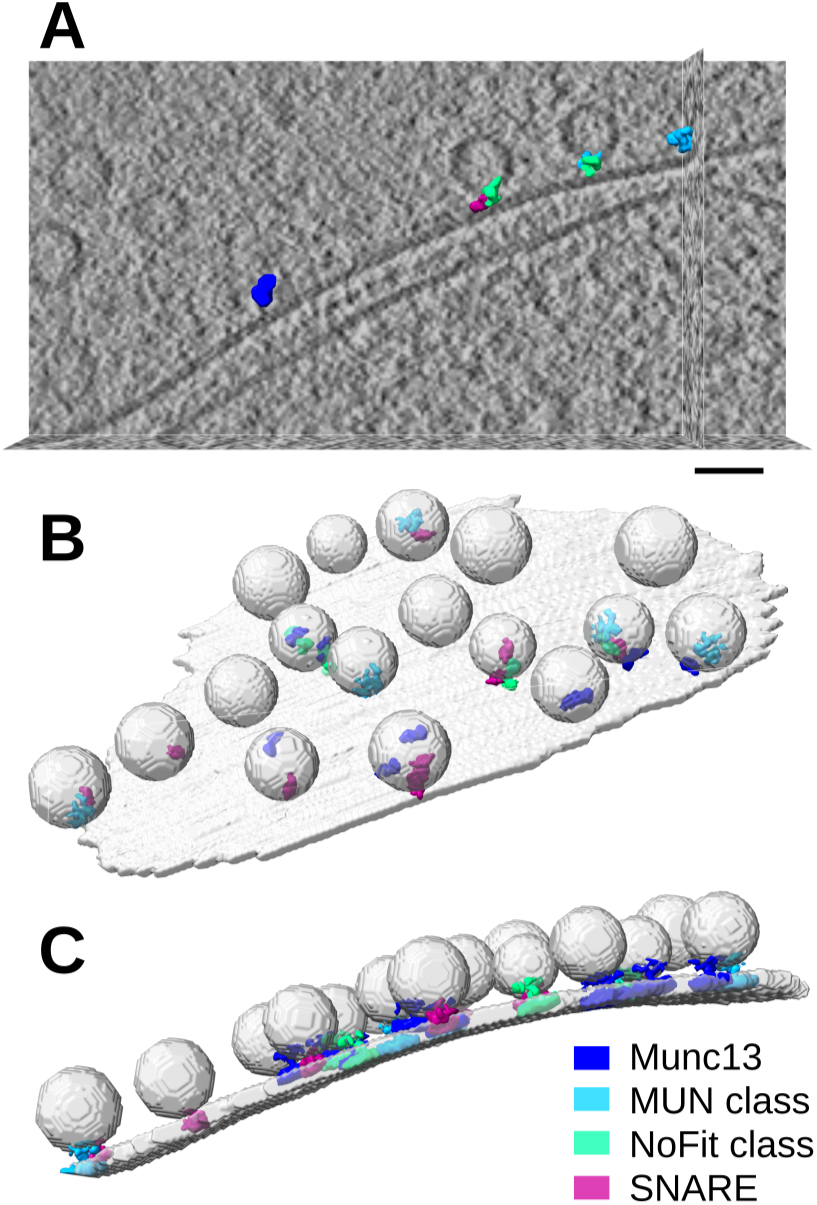
Mapping average densities of Munc13, SNARE complex and two unidentified complexes (MUN and NoFit classes) to positions and orientations of the corresponding tethers in a tomogram. (A) Mapping averages on a tomographic slice. (B) Mapping on the entire tomogram, averages shown contain only cytosolic regions. (C) Mapping on the entire tomogram, averages shown include lipid membrane contributions. Plasma membrane and SVs are shown in light grey. Scale bar 40 nm.

Subtomogram averaging of protein complexes smaller than 500 kDa within their native, cellular environments presents serious difficulties [38]. Recent studies that overcame this limitation include protein bridges between ER and lysosomes (422 kDa), ionotropic glutatmate receptors (400-500 kDa), a ribosome-free translocon complex (∼260 kDa), and molecular bridges between ER and mitochondria that include a genetically encoded fluorophore (214 kDa) [29, 39, 40, 41]. Therefore, the averages we report here of the complex comprising synaptic SNARE proteins, synaptotagmin-1 and complexin-1 (132 kDa) and Munc13 (196 kDa) expand the boundaries of cryo-ET. In addition, the previous subtomogram averages of protein bridges between lipid membranes together with our results further support the notion that neuronal synapses provide a versatile biological system for cellular cryo-ET investigations at the molecular level [42].

Subtomogram averages of short tethers show that they most likely contain the SNARE complex, complexin-1 and synptotagmin-1. This is in agreement with previous results showing that individual short tethers are sufficiently large to accommodate the SNARE complex [33]. Our data fully support the atomic models where the synaptotagmin C_2_B domain is oriented towards the plasma membrane and binds the SNARE four-helix bundle via the primary or tripartite interface, but are not compatible with positioning of the SNARE bundle bound C_2_B domain close to the SV. The primary and the tripartite models radically differ in the positioning of complexin-1, but we could not distinguish between them because they fit into the averages equally well. The current biochemical evidence strongly supports the primary model, but does not completely rule out that the tripartite model exists in vivo [22], leaving the possibility that both conformations we observed are physiologically relevant. Alternatively, and based on the notion that the four-helix SNARE bundle is expected to be at least partially unzippered before the fusion trigger, the C-terminal region of the SNARE four-helix bundle may be more spread out than indicated by the available fully zippered atomic models. Because complexin-1 is located close to the C-terminal side of the four-helix bundle, it is possible that the density occupied by complexin-1 in the tripartite model is actually formed by the unzippered regions of SNARE proteins, which would be consistent with the primary model.

AI classification and subtomogram averaging of intermediate tethers corroborate and extend the results we had obtained by fitting the Munc13 model into a subset of individual tethers directly in tomograms [33]. Our data indicate that a majority of intermediate tethers are composed of Munc13, that they make an angle smaller than 45° to the plasma membrane, and that the DAG binding site and the polybasic region of C_1_ and C_2_B domains face the plasma membrane. This slanted orientation is in agreement with one of the previously proposed models based on biochemical and structural data [19] and observed in cryo-ET studies that imaged a purified and reconstituted portion of Munc13 [43, 21]. However, these studies also proposed that Munc13 can adopt an almost perpendicular orientation to the membrane. Considering that long tethers dominate in the absence of Munc13 [33], and that some of the long tethers (*>*12 nm) may contain Munc13 in the close-to-perpendicular orientation, our data indicate that under physiological conditions, the close-to-perpendicular Munc13 state is less stable and transitions to the more stable, slanted state.

The averages from the two smaller classes, comprising about one third of intermediate tethers, were too small to accommodate the Munc13 model, even though the model comprises only 55% of the entire Munc13 sequence. These tethers were shown before to require the presence of Munc13, while some of them do not contain SNAP-25 [33], indicating that they contain individual SNARE proteins, or other proteins, such as Munc18-1, as proposed before [44, 45]. In essence, the data indicate the existence of a protein complex acting downstream of Munc13 and upstream of SNARE complex formation.

Regarding methodology, several image processing developments were necessary to achieve the present results (Figure S1). First, tethers were detected and localized in an automated, template free manner, not requiring visual detection and using only the information present in the tomograms [35]. In this way, we avoided labor intensive manual picking and picking strategies that oversample membranes thus requiring extensive particle selection [46]

Second, the presence of lipid membranes can dominate subtomogram alignment at the expense of membrane-bound complexes. This is further aggravated by the small size of tethers and the presence of two lipid membranes, located at a variable distance to each other, one being highly curved. Therefore, we designed particle preprocessing approaches and subtomogram averaging improvements that increase the structural information of tethers relative to that of membranes or other protein complexes present in the vicinity of tethers, focus the alignment on tethers to constrain the space of the alignment parameters that is explored during subtomogram averaging, and to extract the cytosolic parts of tether averages. These methods significantly improved averaging of both short and intermediate tethers.

Third, the structural and molecular variability negatively affects subtomogram averaging. We optimized and trained feature based machine and deep learning classifiers, and showed that they generated structurally distinct classes of intermediate tethers, in contrast to the standard image based approach. We used feature based NN classification instead of an image based approach, because the former requires a lower number of inputs and model parameters, and has a lower input redundancy arising from image translation and rotation, which reduces the likelihood of data overfitting.

Fourth, because most proteins present in synapses and other specialized cellular compartments show a sparse, non-periodic organization, a relatively low number of proteins of interest is captured by in situ cryo-ET. To alleviate this problem, we sought to increase the information content per particle in two ways, by incorporating information from tether segmentation and characterization in the preprocessing, and by designing an extensive but non-correlated set of particle features. Indeed, we found that the design and selection of tether features positively influenced classification results. Furthermore, to avoid overfitting in classification, which is especially important for small datasets, we used simple classifier models and carefully optimized them, instead of using complex or pretrained models. In addition, multiple preprocessing approaches and classification methods served to obtain robust outcomes.

In conclusion, the data presented here provide direct evidence for Munc13 and SNARE-dependent complexes that bridge SV and the active zone plasma membranes and place constrains on their orientations and conformations. The image processing methods we developed add to the body of computational tools for structural analysis of synapses and membrane-bound proteins [47, 48], and are applicable to subtomogram averaging of other sparsely distributed membrane-bridging protein complexes imaged by cryo-ET in crowded cellular environments.

## Methods

### Sample preparation and cryo-ET acquisition

Tomographic series used here were acquired in the scope of a previous publication [33]. These show mammalian central nervous system synapses from synaptosomal preparation, which is a well-established model that responds to pre- and postsynaptic stimulation [49, 50, 51, 52, 41], preserves the macromolecular architecture and is an excellent substrate for cryo-ET imaging at the molecular level [31, 53, 23, 54]. In short, mouse E18 or P0 hippocampal organotypic slices were homogenized at DIV 28-30 and centrifuged to yield the P2 (crude synaptosomal fraction). The resusspended fraction was vitrified and imaged under cryo-conditions at 300 kV on Titan Krios (Thermo Fisher) operated in the zero-loss energy filter mode. Synaptosomes were detected in mages and cryo-ET series were acquired from -60° to 60° with a 1.5° - 2° angular increment, on a direct electron detector device (K2 Summit, Gatan) operated in the counting mode, at 0.44 nm pixel size at the specimen level. Volta phase-plate [55] with nominal defocus of 0.5-1 μm was used and the total dose was kept *<*100 e^−^/^°^A^2^. The series were aquired using SerialEM [56] and motion corrected using MotionCor2 [57]. All aligned tilt series and tomograms were deposited at the Electron Microscopy Public Image Archive (EMPIAR-12512). All tomograms deemed technically and biologically acceptable in the previous study ([33]) were used here. In short, tomograms were used if they did not contain any signs of ice crystal formation such as ice reflections or faceted membranes, there were no flaws in the acquisition, and the synapses did not show signs of deterioration such as elliptical small vesicles or strong endocytotic features.

Here, 77 tomograms were reconstructed by weighted back projection using Imod [58] at 0.88 nm pixel size (as opposed to 1.76 nm in the previous study).

### Membrane segmentation, tether detection and analysis

These tasks were performed in the scope of a previous publication [33]. In short, active zone plasma membrane and SVs were manually segmented. Tethers were detected, localized and segmented by hierarchical connectivity procedure implemented in Pyto package and analyzed using Pyto package [30, 35]. This tether detection procedure is template-free and does not require visual detection, as it automatically finds bridges between the pre-segmented SVs and the plasma membrane at different greyscale levels. Tether analysis included the determination of their morphology, greyscale, topology, and distance to other structures.

### Tether annotation, feature extraction and selection

Tethers were annotated manually by rigid body fitting of the atomic model of Munc13 comprising C_1_, C_2_B and MUN domains [[37] PDB id: 5ue8] (Figure S8 B) into individual tethers from randomly selected tomograms, using UCSF ChimearX software [59]. Tethers that could accommodate the Munc13 model were assigned to the Munc13 class, those that could not accommodate the MUN domain but not the entire Munc13 model to the MUN class, and the remaining tethers to the NoFit class. The total of 437 tethers were annotated, resulting in 46, 66 and 325 tethers, in the NoFit, MUN and Munc13 classes, respectively.

The features obtained from our previous tether characterization include morphological, greyscale density and topological properties of tethers, as well as the minimal tether length that approximates tether shape and the minimal distance between the tether-associated SVs and the plasma membrane [35, 33]. Here we further analyzed the previous data to obtain different modalities of tether length arising from the tether shape and the extended nature of tether-membrane contacts (minimal, median and maximal) and features related to the threshold at which tethers were detected. Together, these defined 14 Basic features.

The Derived features (11 in total) were obtained from the Basic features. Some of these provide normalization by tether volume, thus reducing the influence of tether size, such as the surface to volume ratio and the ratio of topologically distinct loops and volume. Others combine different tether length features, such as relative difference between maximal and minimal tether lengths, which provides information about the extension of the membrane-bound tether region, and the ratio between tether lengths determined with and without considering tether shape, which provides the tether curvature estimate. Most of the Derived features use non-linear Basic feature combinations, because they may improve classification, especially when linear machine learning approaches are used.

Plotting pairwise distributions of features from the Basic and Derived features for all tethers, and computing univariate statistics based on ANOVA and Mutual information measures for annotated tethers allowed us to discard highly correlated and low score features resulting in 16 “Selected” features (Figures S5, S6). Different feature sets were generated by (i) optional logarithmic transformation of highly skewed features (previously converted to positive values *>*1 if needed) (ii) standard (normal) or robust scaling. These resulted in robust, standard, logrobust and logstd scaled feature sets.

### Tether classification by machine learning (excluding deep learning)

We evaluated the following machine learning models (excluding deep learning): Logistic regression with L2 regularization, with linear and 2nd order polynomial feature combinations, SVM classifier with the Gaussian radial basis function kernel, Decision trees, Gradient boosted trees and Ada boosted Logistic regression, all implemented in Scikits-learn [60]. The hyperparameter search was performed by cross-validation over large range of values (parameter 10^−3^ to 10^3^ for parameter *C* of LogReg and SVM, 10^−2^ to 10^2^ for *γ* of SVM). Features of 437 annotated tethers were randomly separated into training (327) and testing sets (110 tethers). Cross-validation with F_1_ macro scoring function was used for hyperparameter search. Models were evaluated using F_1_ macro score on test data, but we also calculated accuracy and weighted F1 score, which applies a higher weight to classes with more elements. For completeness, evaluation was also performed on the training set.

The optimized LogReg (*C* = 1) and SVN models (*C* = 100 and *γ* = 0.001) were trained on the entire annotated dataset and used to classify intermediate tethers, resulting in 417, 201 and 142 particles in Munc13, MUN and NoFit classes, respectively, by LogReg and 425, 183 and 152 particles by SVM classifier.

### Tether classification by deep learning

As above, annotated tethers were randomly separated into training (327) and testing sets (110 tethers). We used NNs comprised of fully connected layers as implemented in PyTorch. Hyperparameters and feature sets were optimized using a series of cross-validations on the training set by first varying multiple hyperparameters at the same time, and then by varying a single hyperparameter at a time starting from the optimized network. The optimal network contained four 18-node hidden layers (the first and the last layer had to equal the number of features and the number of classes, respectively), the weights i initialization was Kaiming normal, and we used batch normalization, dropout rate of 0.3, Adam optimizer, ReLU activation function and F_1_ macro loss function. Batch size was 30, the learning rate for the first 1000 epochs was 0.001 and 0.0003 for epochs 1000 - 2000.

Regarding hyperparameter optimization, the choice of the optimizer was very important, with Adam largely overperforming stochastic gradient descent and AdaGrad optimizers (F_1_macro 0.37 and 0.40, respectively). Batch normalization and optimization of the dropout rate provided significant improvement of F_1_ macro score (0.70 - 0.74) and decreased overfitting. While ReLU activation function was optimal, ELU and leaky ReLU provided slightly decreased F_1_ macro scores (0.76 in both cases). We also optimized the learning rate schedule and the number of training epochs.

Models were evaluated using F_1_ macro score on test data. For completeness, evaluation was performed with F_1_ macro, F_1_ weighted and accuracy evaluation scores. To be able to also use F_1_macro as the loss function, we implemented a smooth (differentiable) version of F_β_ metric, which takes class probabilities generated by the last network layer to calculate first the confusion matrix and then different F_β_ scores.

The optimized model was then used to predict classes for non-annotated intermediate tethers, resulting in 441, 229 and 90 particles in the Munc13, MUN and NoFit classes, respectively.

### Particle preprocessing

Segmented SV and plasma membranes, and tethers detected by the template-free hierarchical connectivity method were obtained previously from 77 tomograms [35, 33]. These were used here to determine tether contact coordinates by finding the center-of-mass coordinates of the tether region that is in direct contact to the plasma membrane (direct contact was defined as pixels sharing a face), as well as the two Euler angles that define the membrane normal direction [29] (Figure 1B). We then determined tether centers as points located 10 pixels from the tether contacts, along the membrane normals, in the cytoplasmic direction. The tether centers were used as centers for all subtomograms that were subsequently extracted, as explained below. Furthermore, initial tether alignment angles were set using the membrane normal angles. In this way, transforming subtomograms according to the initial alignment parameters aligns membrane and the tether contacts of all particles.

The Plain tether particle set was obtained by extracting subtomograms from the previously obtained tomograms, the Borders particle set from the segmented SV and plasma membranes, and the tether mask set from the segmented tethers. Consequently, each tether particle had corresponding borders and tether mask particles, all three aligned with each other. The box length was 64 pixels in all cases (pixel size was 0.88 nm).

Initially, additional tether particle sets were generated as follows. The filtered SV tether particle sets were obtained by low-pass Gaussian filtering the region corresponding to SVs, as defined by the boundaries set of the Plain tether set, at cutoff frequencies between 1/0.5 to 1/5 nm. Similarly, filtered SV & PM tether sets were obtained by low-pass Gaussian filtering region corresponding to SVs and plasma membranes (independently from each other), as defined by the boundaries set, of the Plain tether set at cutoff frequencies between 1/0.5 to 1/5 nm. In all cases (Plain, filtered SV and filtered SV&PM sets). The focused particle sets were obtained by randomizing all pixels outside the tether mask, with or without morphological dilation of 5 pixels, in all directions or only over the cytosolic region.

After a visual assessment of averages generated from the above particle sets, the following four final tether particle sets were chosen: (i) Plain, (ii) Filtered SV using 1/5 nm cutoff, (iii) Filtered SV&PM using 1/5 nm and 1/0.5 nm cutoffs for SVs and plasma membrane, respectively and (iv) Focused, using 5 pixel dilation over the cytosol with Filtered SV&PM superimposed on the SV and plasma membrane regions (Figure S2).

For subtomogram averaging, pixels outside of cylinders of radius 10/15 and height 22/30 pixels (short / intermediate tethers), starting 10 pix away from tether contacts in the extra-cellular direction and having axes aligned with the membrane normals were randomized, as we and others have done previously [41, 61].

Relion format particle star files were automatically generated for each particle set.

### Subtomogram averaging of tethers

Subtomogram averaging, comprising refinement and optionally 3D classification, was performed in Relion 3.0 [62].

The three classes obtained by the (image-based) Conventional3D class averages were evaluated based on the class averages size and associated to Munc13 (the largest), MUN and NoFit (the smallest) classes. The resulting Munc13 classes contained 186 in Plain, 359 in Filtered SV, and *<*100 particles in the other two preprocessing cases.

All refinements were performed de novo, that is without the use of external references, and without any symmetry. Initial references were obtained by aligning and averaging all particles according to the two angles determined from membrane normals and randomizing the third angle (around the normal direction) to remove the missing wedge. During the refinement, particle half-datasets were processed independently and the final resolution was determined by Fourier shell correlation at the 0.143 level according to the “gold-standard” procedure [63], as implemented in Relion. Solvent masks, matching the preprocessing mask sizes, were applied during refinements to ensure unbiased resolution estimates. For postprocessing, including FSC and resolution determination, we used soft cylindrical masks having 10 pixel radius and 14 pix height for short and 15 pixel radius and 20_pix height for intermediate tethers. Box size was 64 pixel and pixel size was 0.88 nm.

All refinements used the constrained alignment, which allows only small changes of angles corresponding to the direction of the membrane normal vector and small spatial displacements. The alignment around the third angle (around the normal vector) is optimized over the entire angular range. To implement this, we set the prior values for angles *tilt* and *psi* in Relion particle star files to the two angles defining the normals to the membrane, and specified small values (3.66) for the standard deviations of these two angles in the refine command options.

Short tethers (321 in total) were detected, extracted, preprocessed and averaged as explained above. Intermediate tethers (760 in total) were processed in the same way except that they were classified by Relion, or by LogReg, SVM or NN classifier, before averaging. Conventional3D classification into three classes was done using Relion 3D classification of each preprocessed set separately, resulting in 186, 351 and 223 particles for Plain, 359, 261 and 140 for Filtered SV, 80, 392 and 288 for Filtered SV&PM, and 87, 155 and 518 for Focused preprocessing, where classes were ordered by their similarity with the Munc13 model. AI classification resulted in 417, 201 and 142 particles for LogReg, 425, 183 and 152 for SVM and 441, 229 and 90 for NN classifier, in Munc13, MUN and NoFit classes, respectively.

### Determination of the cytosolic region of tether averages

Subtomogram averaging of the boundaries particle set (segmented plasma membrane and SVs) was obtained by simple averaging using the final alignment parameters obtained from tether subtomogram averaging, This procedure was repeated for all tether averages. In this way, each tether average was assigned and perfectly aligned with a corresponding boundary average. Because of the initial tether particles orientation and the constrained alignment, all tether and boundary averages were oriented so that plasma membrane was parallel to z-slices.

All boundary averages were independently thresholded to separate plasma membranes and SVs from cytosolic regions, as follows. The x and y coordinates of SV nadir (the point closest to the plasma membrane) in boundary averages were determined by visual inspection of z-slices. Density traces perpendicular to the plasma membranes (along z-axes), of 5×5 pixel region centered on SV nadirs showed locations of the plasma membrane and SVs along z-axis, and were used to determine the density level at which the plasma membrane was 8 pixels thick (Figure S4C, S10). These density values were used as thresholds to generate boundary masks. In addition, the position of the cytosolic face of the plasma membrane was determined as the higher limit of the plasma membrane location along z-axis. To allow easier comparison between different cases, the traces were shifted so that the threshold density value and the z-coordinate of the cytosolic plasma membrane face were both 0.

For visualization of tethers, boundary masks were imposed on the corresponding tether averages to remove the membrane contributions without modifying the cytosolic portions. The visualization threshold was set so that the membranes in tether and boundaries averages agree. While this already resulted in all short tether averageshaving similar volume, the thresholds were finely adjusted so that the cytosolic tether volumes were between 290 and 300 nm^3^. In addition, all disconnected parts of tether averages were kept in Figures S3, S8 and S11, and only the largest parts are shown in Figures 2 and 3. Visualization was performed using UCSF ChimearX software [59].

### Fitting

Fitting was performed using manual rigid body fitting in UCSF ChimearX software [59].

Three different models were fit into the short tether averages. (i) The primary model was created by combining all SNARE helices and synaptotagmin-1 C_2_B domain from PDB id: 5kj7 [64] with SNARE helices (syntaxin and SNAP25) and complexin-1 helix from PDB id: 1kil [65]. (ii) The tripartite model was created by combining the SNARE primary model (without synaptotagmin-1 C_2_B domain) with the synaptotagmin-1 C_2_B domain from PDB id: 5w5c [12]. (iii) The model containing two synaptotagmin-1 C_2_B domains (primary and tripartite) was created by combining the primary model and the tripartite models. In this way, all SNARE models have the same SNARE and complexin helices and differ only in the positioning of synaptotagmin-1 C_2_B domain (Figure S3B). In all cases, synaptotagmin-1 C_2_B polybasic residues (322, 324, 325 and 326) were labeled in blue [11]. In all cases, the priority was to fit the SNARE four-helix bundle and the synaptotagmin C_2_B domain. All three models were oriented so that the polybasic residues were oriented towards the plasma membrane (Figure S3A). In addition, the tripartite model was also fit so that the polybasic domain was facing the SV, resulting in four different fitting modes. To assure a proper comparison, the visualization thresholds of all average densities were adjusted so that their volume was essentially the same (290 - 300 nm^3^).

Two models were used for fitting into intermediate tethers. (i) The Munc13 model comprises Munc13 C_1_, C_2_B and MUN domains and covering 55% of the entire Munc13 sequence PDB id: 5ue8 [37] (Figure S8B). (ii) The MUN model contained only the MUN domain of the Munc13 model. Polybasic residues of C_1_ and C_2_B domains (592, 720, 722, 724, 750) were labeled in blue and the W588 residue at the DAG binding site in purple [66]. In all cases, the polybasic residues and W588 were oriented towards the plasma membrane (Figure S8). Munc13 model was fit into the Munc13 class averages, MUN model was fitted and Munc13 model was attempted to fit into the MUN class averages and MUN was attempted to fit into the NoFit class averages.

### Mapping of average densities

Average densities of Munc13, MUN and NoFit classes of intermediate tethers obtained using Filtered SV&PM preprocessing and NN classification, and of short tethers obtained using Focused preprocessing were mapped in 3D on the exact positions and according to the angular orientation of individual particles, as determined by the outcome of the refinement procedure. Average densities that include the contribution from lipid membranes and also those containing only cytosolic regions were used. The positioning and the rotation of the average densities was performed automatically by Pyto software [35], and ChimeraX [59] was used for visualization.

## Data availability

All data needed to evaluate the conclusions in the paper are present in the paper and the Supplementary Materials. Subtomogram averages of short tethers with Focused preprocessing (Fig 2A, B) and of NN classified Munc13 class intermediate tethers with Filtered SV&PM preprocessing (Fig. 3A) are deposited at the EMDB [67] (EMD-52342, EMD-52343). All aligned tilt series and tomograms generated for a previous study [33] and used here are deposited at the Electron Microscopy Public Image Archive [68] (EMPIAR-12512). Representative tomograms were deposited previously at the EMDB (EMD 16083, 16084, 16085). All other data are available upon request.

## Code availability

All software developed for this work was written in Python 3 language, using classes and functions implemented in Pyto package (version 1.10, available on GitHub https://github.com/vladanl/Pyto) [35], which uses NumPy and SciPy scientific computing packages [69, 70]. In addition, Pandas package is used for particle preprocessing, feature determination and all data manipulation procedures, Scikits-learn and Seaborn for pairwise and univariate feature analysis, Scikits-learn and PyTorch for building machine and deep learning classifiers, and Matplotlib for plotting [71, 60, 72, 73, 74].

## Acknowledgments

This work was supported by the HFSP RGP0020/2019 grant. We would like to thank Wolfgang Baumeister and Juergen Plitzko for their support, and Gabriela J. Greif for critical reading of the manuscript.

## Author information

### Contributions

D.H.O.-B. and C.P. generated data; D.H.O.-B., C.P. and V.L. analyzed data; N.B. provided critical expertise and assisted in result interpretation. V.L. designed and supervised research, wrote software and acquired funding. V.L. wrote the manuscript; all authors edited the manuscript.

### Current affiliations

D.H.O.-B. present address: National Institutes of Health (NIH) 9000 Rockville Pike Bethesda, Maryland 20892, USA

C.P. present address: Neurobiology Division, MRC Laboratory of Molecular Biology, Cambridge Biomedical Campus, Cambridge CB2 0QH, U.K.

### Corresponding author

Correspondence and requests for materials should be addressed to VL.

## Supplementary information

The following supplementary Information is available for this paper: Table S1, Figures S2-S11

## Supplementary Material

**Figure S1:**
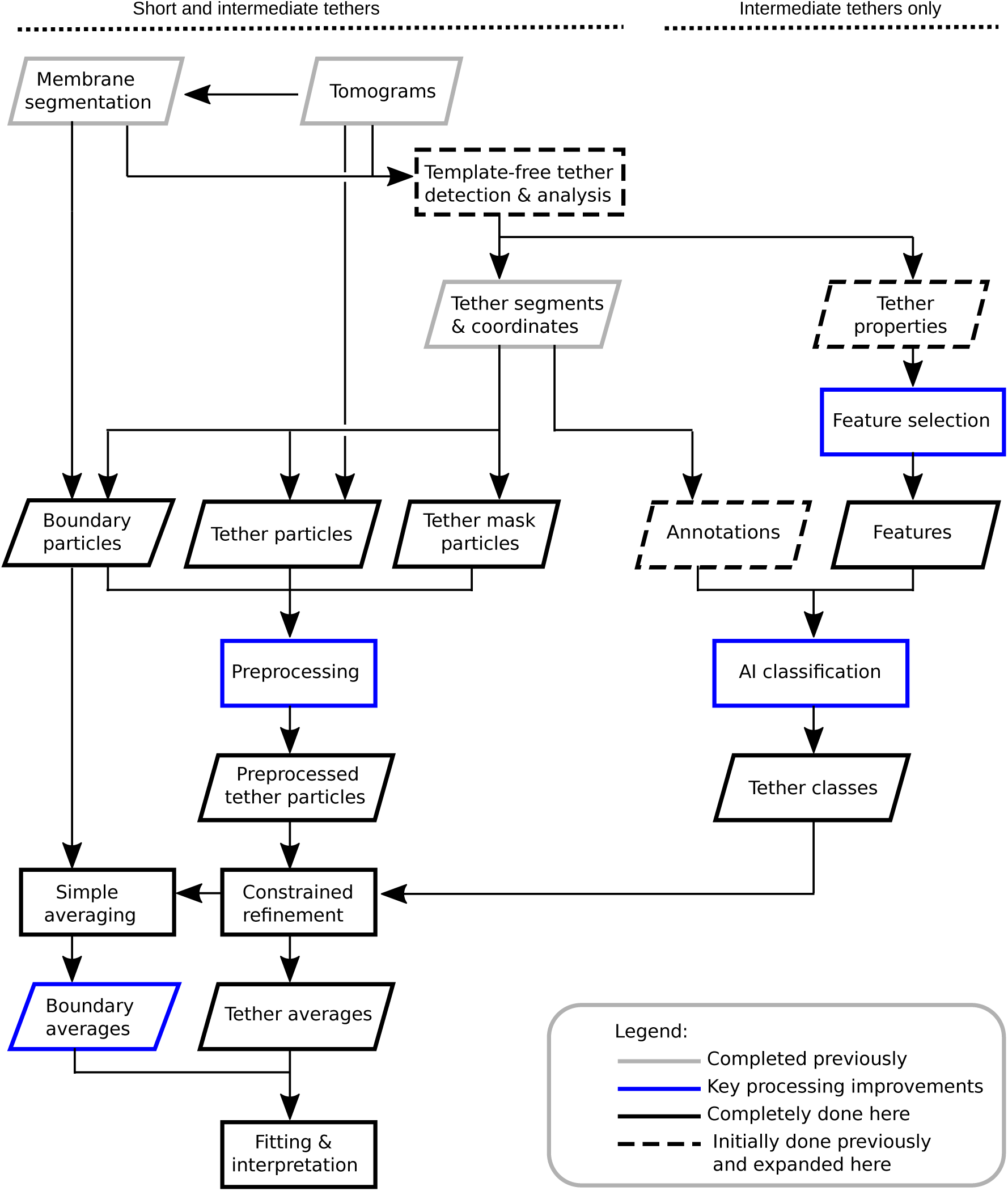
Processing workflow. Data is shown as parallelograms and the processing steps as rectangles. Steps completed in our previous publication [33] are shown in grey, the key processing developments introduced here in blue, other steps completely done here in black, and the steps initially done in the previous but expanded in this publication are indicated by dashed lines.

**Figure S2:**
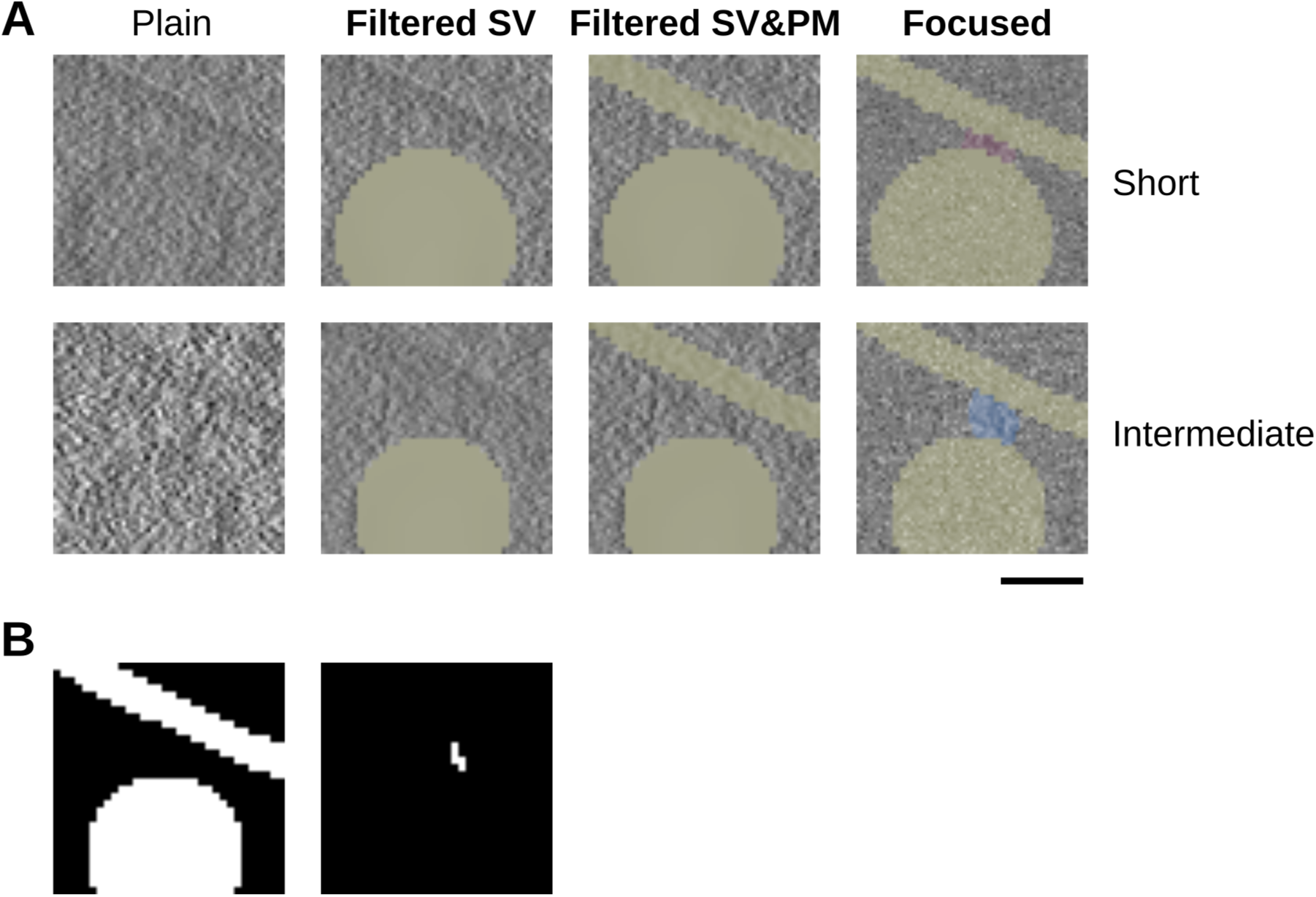
Tomographic slices of particle images. (A) Tether particles, a short tether is shown in the upper row and an intermediate in the lower row. The four preprocessing cases are shown in the columns as indicated, those introduced here are shown in bold. (B) Boundary particle (left) and segmented tether (right). All images correspond to the same tether, slice thickness 0.88 nm. Scale bar 20 nm.

**Figure S3:**
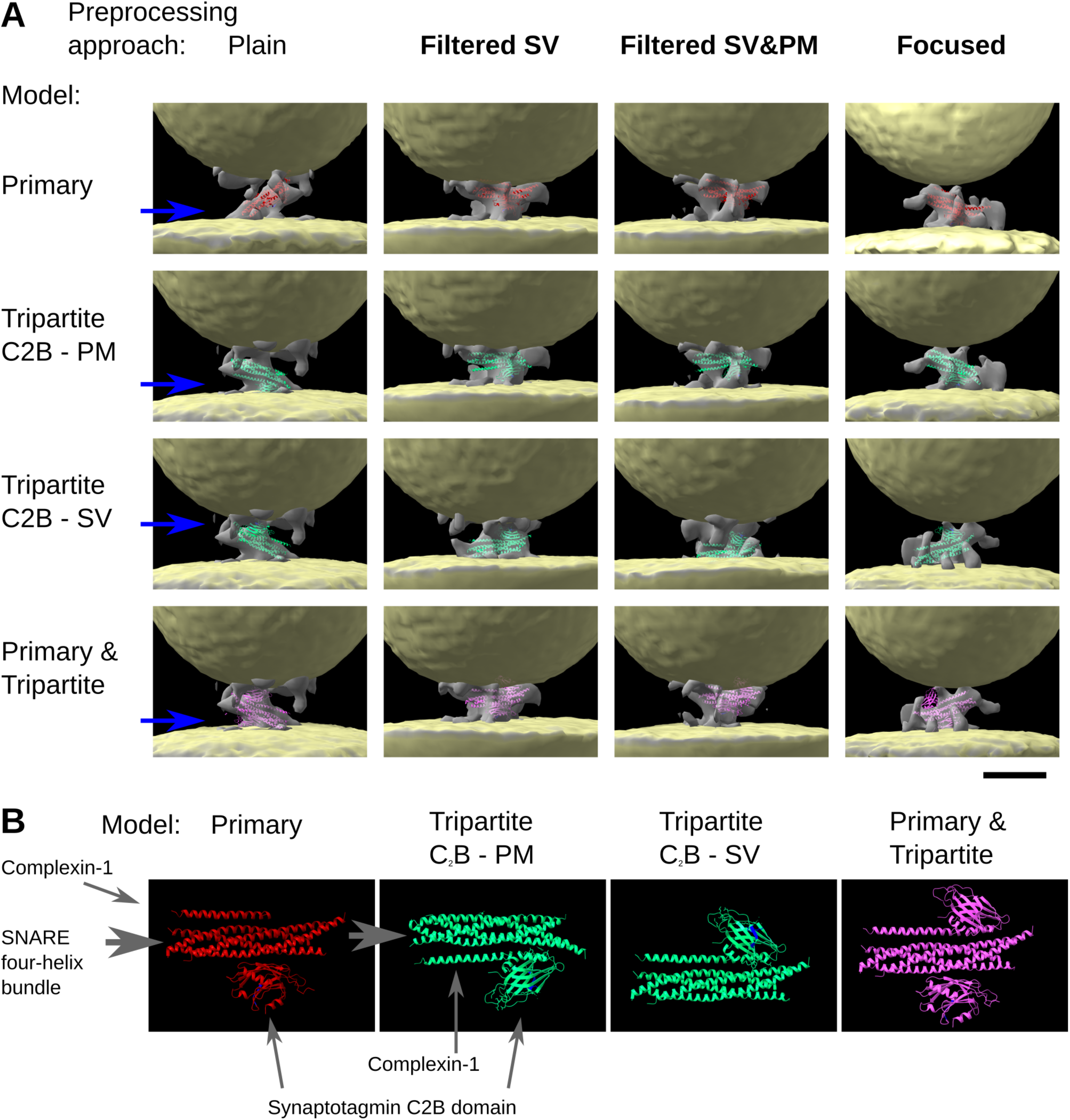
Short tether de novo averages. (A) Averages shown are for all four preprocessing approaches and four models, as indicated. Preprocessings introduced here are shown in bold. Blue arrows points to the approximal height where the polybasic residues of the synaptotagmin C_2_B domain are located. Scale bar 10 nm, images drawn to scale (B) Atomic models, as indicated, drawn to scale. Pynaptotagmin C_2_B polybasic residues are shown in blue.

**Figure S4:**
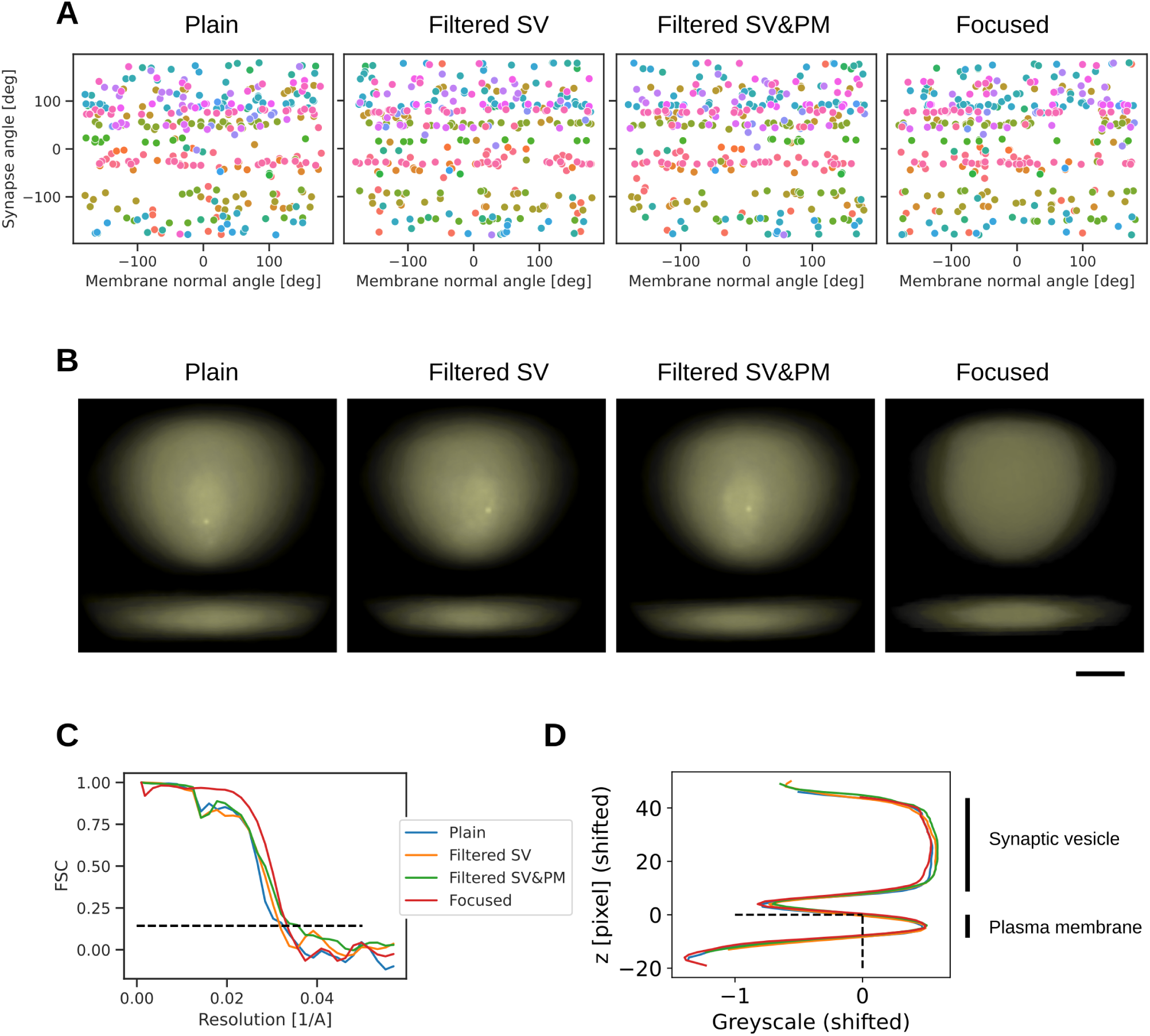
Supporting data for short tether de novo averages. (A) Particle angular distribution. (B) Regions averages, volume rendered. Scale bar 10 nm. (C) FSC curves. (D) Boundaries average density traces. The horizontal dashed line shows the position of the cytoplasmic face of the plasma membrane. The vertical dashed line shows the greyscale level that separates the membranes (*<*0) and the cytoplasm (*>*0). (C) and (D) share the same legend.

**Figure S5:**
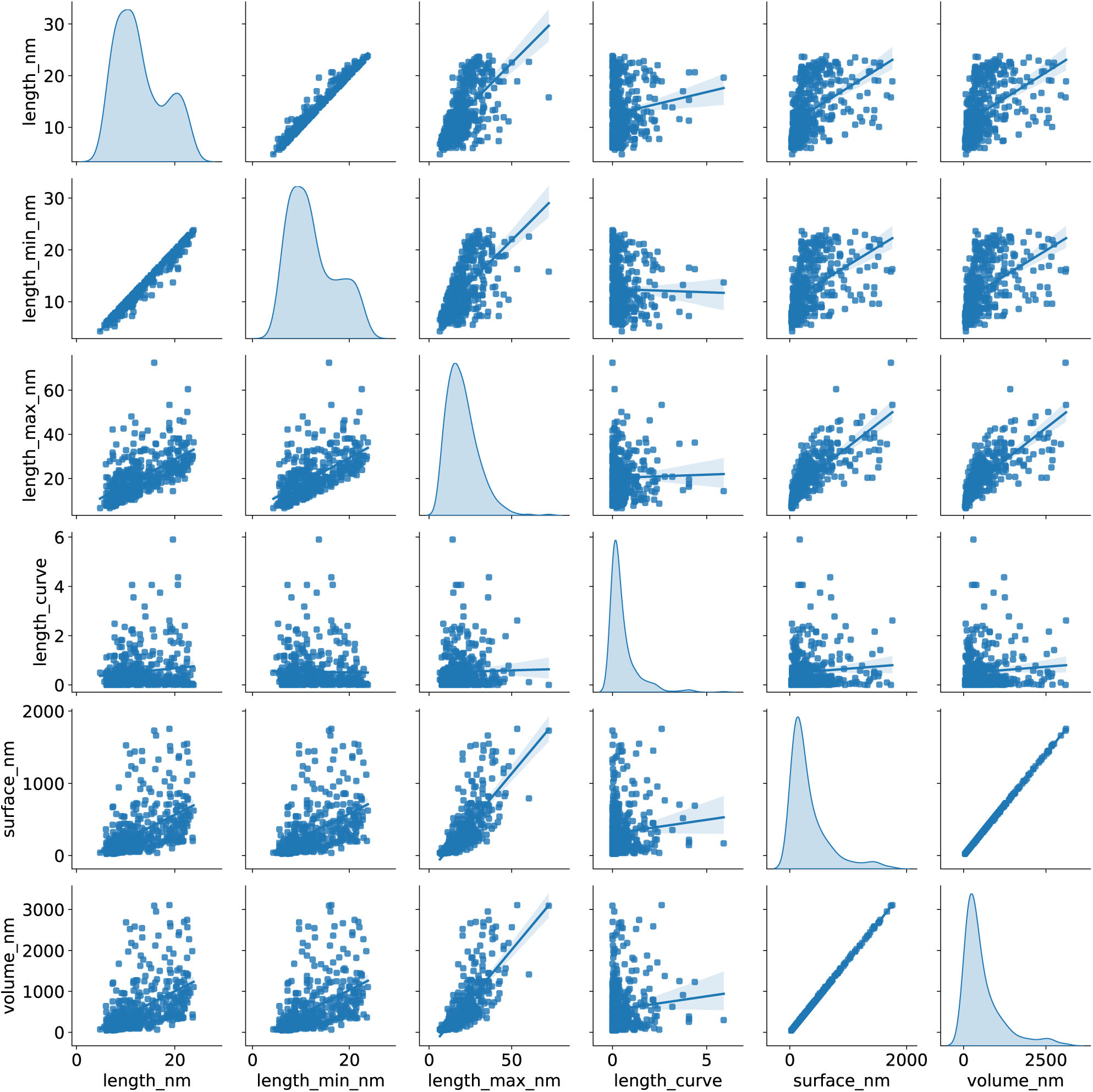
Example of removing correlated tether features. Features shown are (in this order): minimum length with curvature approximation, minimal straight length, maximal straight length, difference between length with curvature approximation and straight length, surface, volume. Minimal straight length and surface were removed.

**Figure S6:**
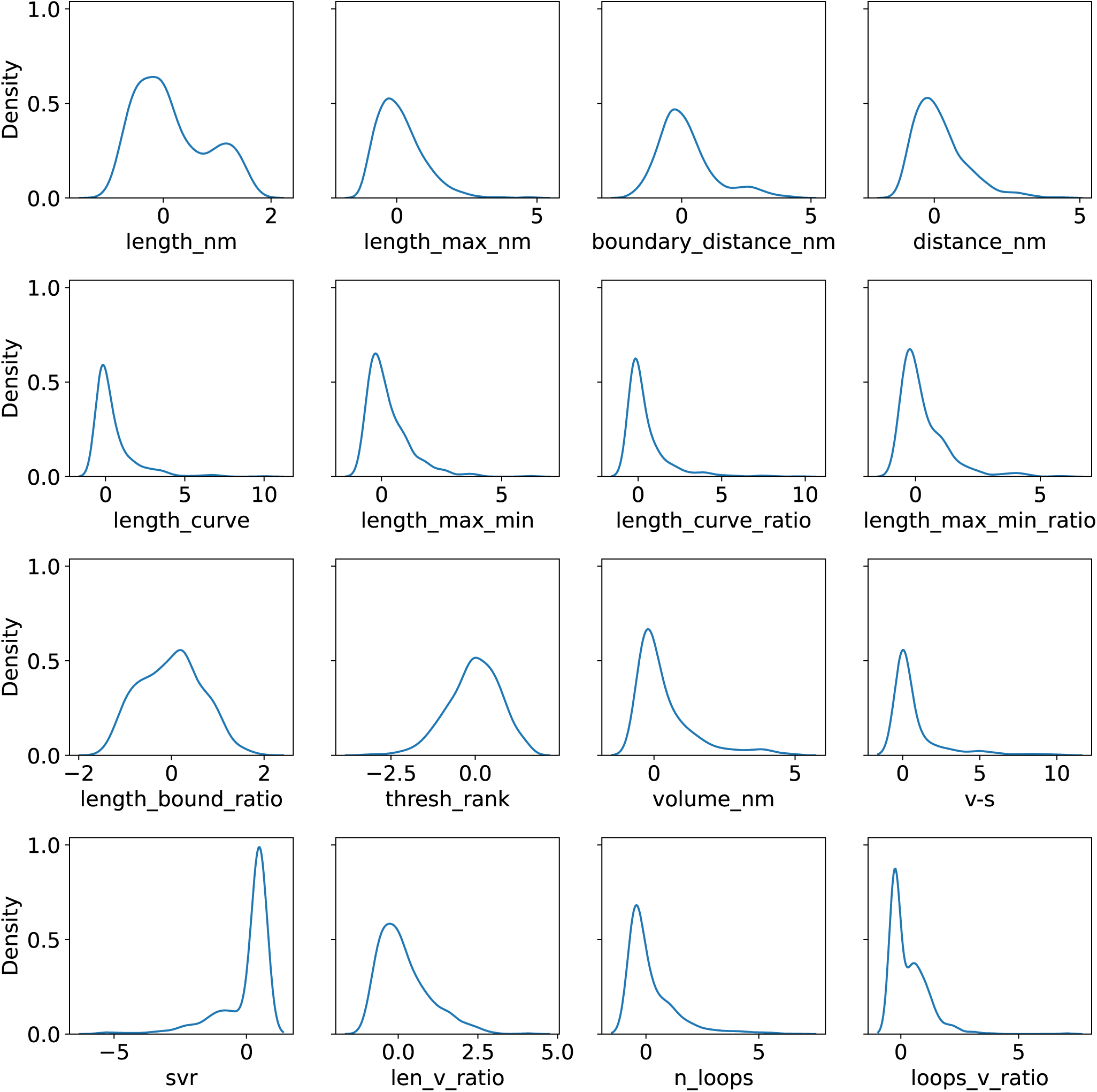
Univariate distribution of the Selected features.

**Table S1:**
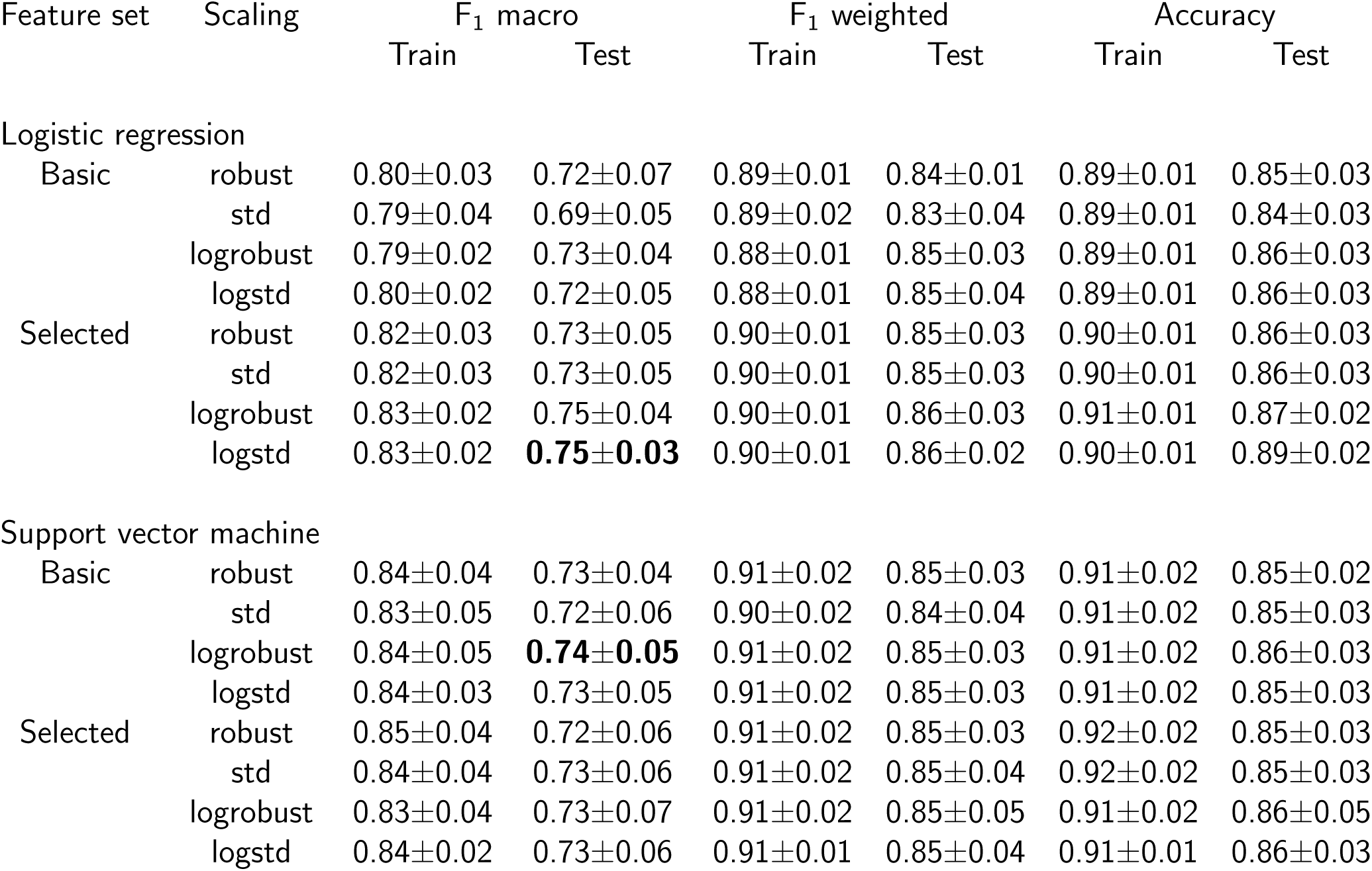
Logical regression cross-validation scores. The training and test scores were obtained for 22 random training-test splits (mean ± std). For each combination of the feature set and scaling, the scores shown were obtained for the values hiperparameter *C* identified by cross-validation using the F_1_ macro metric.

**Figure S7:**
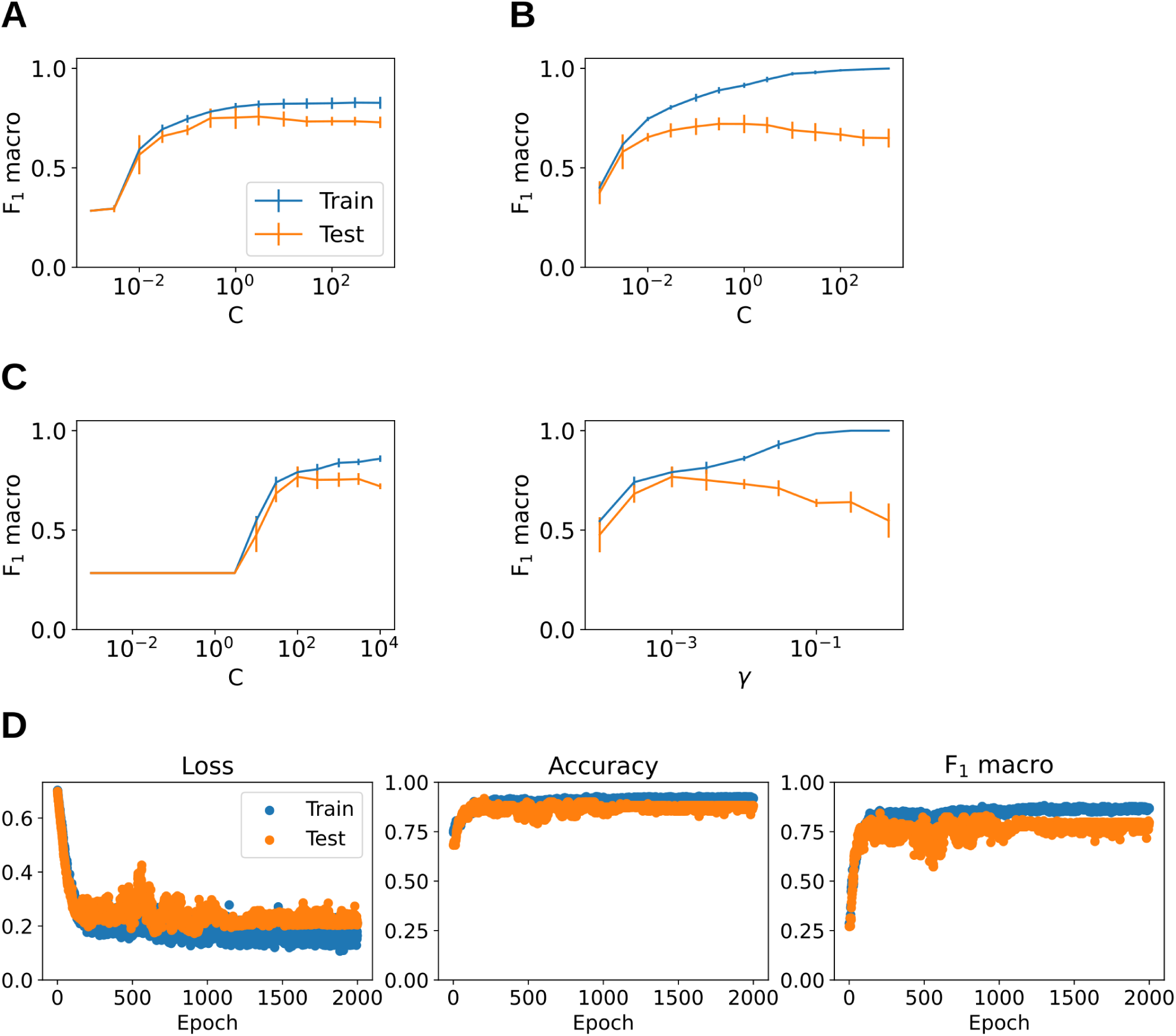
AI classification validation graphs. (A) Logreg linear (B) Logreg polynomial. (C) SVM for γ=0.001 (left) and C=100 (right). (D) NN score graphs for the best performing model. Selected features and logstd scaling were used.

**Figure S8:**
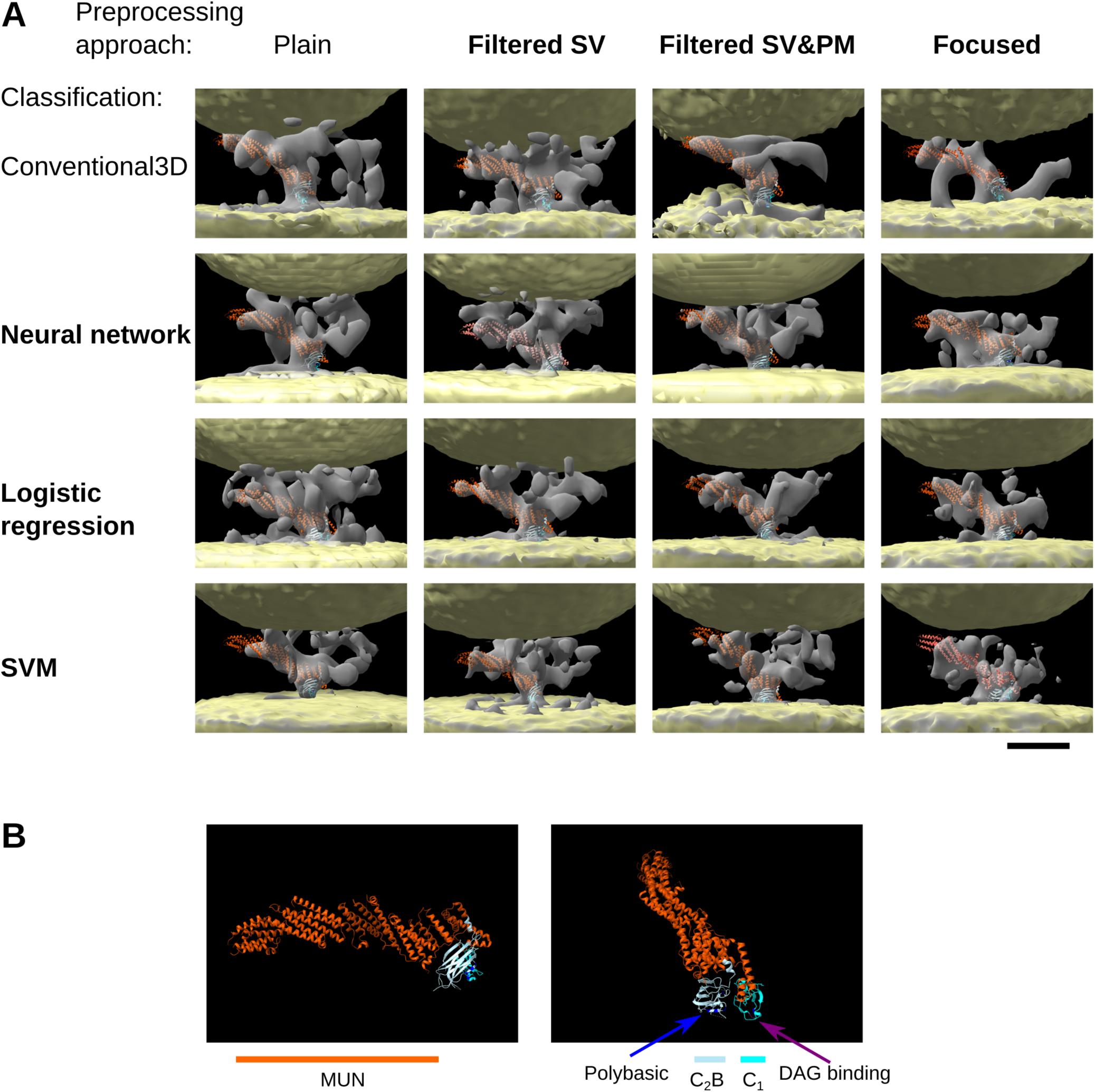
Intermediate tether de novo Munc13 class averages. (A) Averages shown are for all four preprocessing approaches and and all classifications, as indicated, those introduced here are shown in bold. Average density (grey), vesicle and plasma membranes (yellow). Scale bar 10 nm. (B) Domains of Munc13 atomic model shown in different orientations. Munc13: C_1_domain (cyan), C_2_B domain (light blue), MUN domain (orange), polybasic residues on C_1_ and C_2_B (blue), W588 at the DAG binding site on C_1_ (purple).

**Figure S9:**
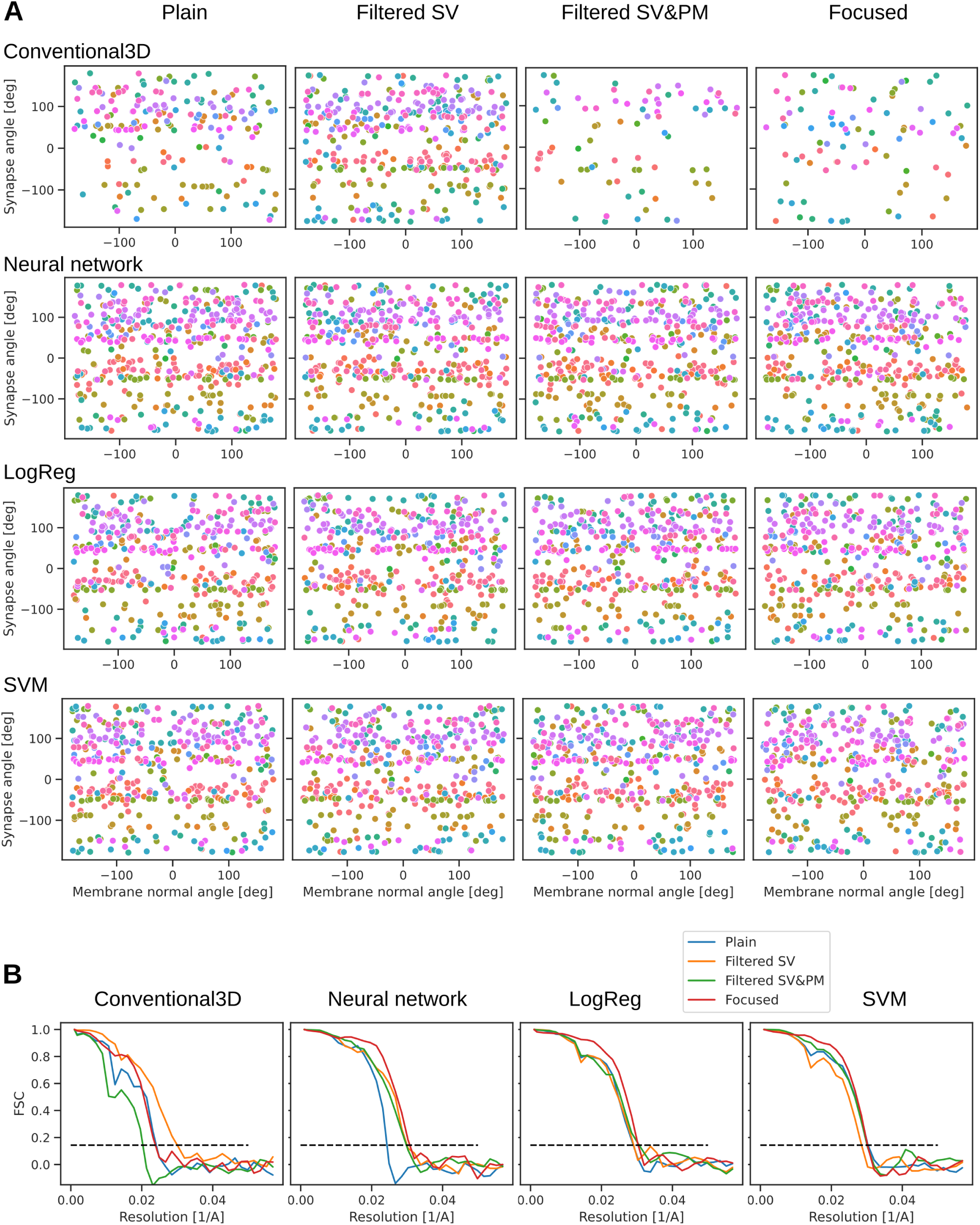
Supporting data for intermediate tether de novo Munc13 class averages. (A) Particle angular distribution. (B) FSC curves. Data shown for all preprocessing cases and classification methods.

**Figure S10:**
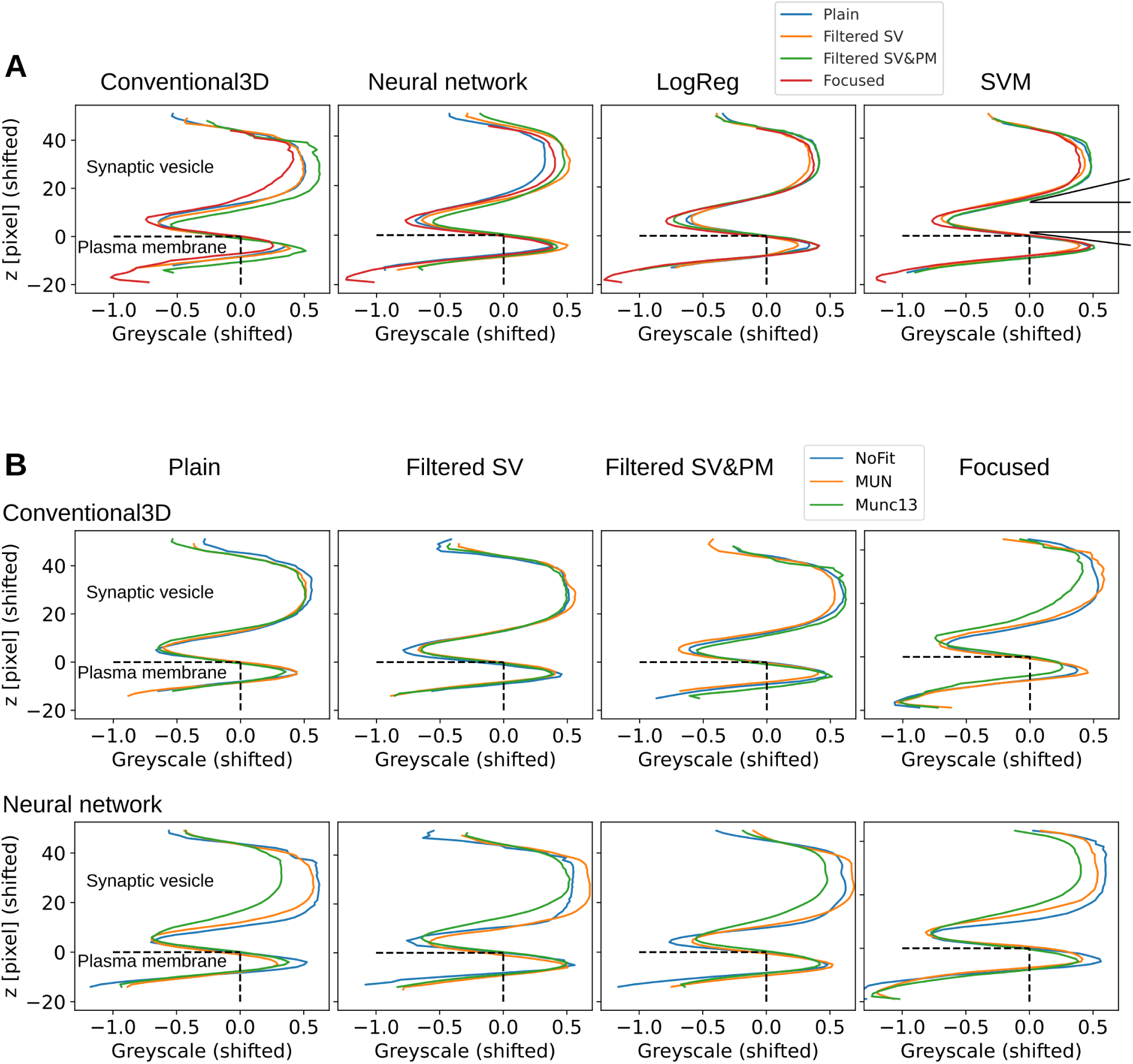
Intermediate boundaries average density traces. (A) Munc13 class traces shown for each classification method separately. (B) Traces for all classes of Conventional3D and NN classification, shown for each preprocessing approach separately. Horizontal dashed line show the position of the cytoplasmic face of plasma membranes. Vertical dashed lines show the greyscale level that separates the membranes (*<*0) and the cytoplasm (*>*0). The two angles at the rightmost graph show that the absolute value of the slope is higher at the SV than at the plasma membrane.

**Figure S11:**
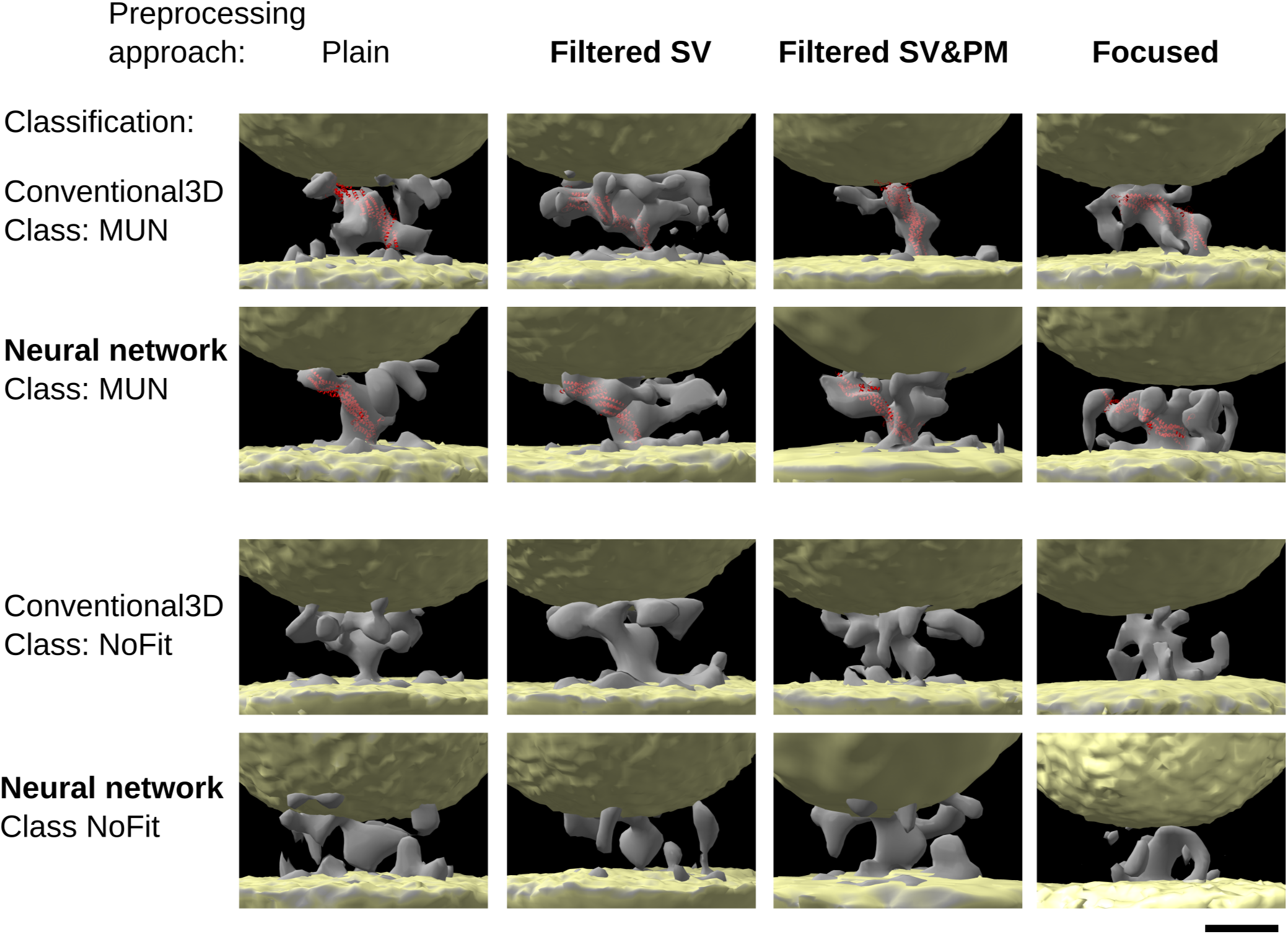
Intermediate tether de novo MUN and NoFit class averages. Averages shown are for all four preprocessing cases, and for Conventional3D and NN classifications, as indicated, those introduced here are shown in bold. (A) MUN class averages. (B) NoFit class averages. Average density (grey), vesicle and plasma membranes (yellow), MUN domain model (red). Scale bar 10 nm.

